# Cell-type-resolved DNA methylation profiling in the cortex reveals female-specific X chromosome differences in autism

**DOI:** 10.64898/2026.09.16.749446

**Authors:** Alice Franklin, Jonathan P Davies, Barry Chioza, Joe Burrage, Emma M Walker, Rosemary Bamford, Ann C Babtie, Georgina E.T. Blake, The APEX Consortium, Simon Baron-Cohen, Emma L Dempster, Eilis Hannon, Jonathan Mill

## Abstract

Sex differences are a prominent feature of many neurodevelopmental conditions, particularly autism for which approximately three males are diagnosed for every female. Although epigenetic dysregulation has been implicated in autism, the extent to which autism-associated epigenetic differences vary by sex and neural cell type remains poorly understood. Using fluorescence-activated nuclei sorting (FANS), we isolated neuron-enriched (NeuN+), oligodendrocyte-enriched (SOX10+), microglia-enriched (IRF8+) and astrocyte-enriched (NeuN-/SOX10-/IRF8-) nuclei from post-mortem prefrontal cortex tissue from 24 autistic (70.8% male) and 23 non-autistic (69.6% male) donors, and performed cell-type-resolved epigenome-wide association studies (EWASs) of DNA methylation. We identified cell-type-specific autism-associated differentially methylated positions (DMPs), with microglia harbouring the largest number of associations. Modelling the interaction between sex and diagnosis revealed a striking female-specific signature concentrated on the X chromosome in microglia. At the majority of these sex-by-autism DMPs, autistic females showed lower DNA methylation compared to control females consistent with an altered X chromosome inactivation (XCI) profile. Across the X chromosome, autism-associated DNA methylation differences were significantly larger in females than males and were greater at sites annotated to genes normally subject to XCI than at genes that escape XCI. Together, these data further support the importance of microglia in autism and suggest that altered X chromosome dosage regulation may contribute to sex-dependent molecular mechanisms in autism.

## INTRODUCTION

Autism is a highly heritable [1,2] neurodevelopmental condition marked by differences in social interaction, communication and unusually restrictive interests and repetitive behaviours [3]. A striking feature of autism is the marked sex difference in prevalence with approximately three males diagnosed for every female [4]. Whilst sex differences in clinical presentation and diagnostic ascertainment likely contribute in part to the sex bias, a substantial male bias remains after accounting for these factors, implying that biological mechanisms may also contribute to these differences [4]. One hypothesis for the male bias in autism proposes that females require a greater genetic load of autism-associated variants to pass the diagnostic threshold [5]. This “female protective effect” (FPE) is supported by the observation that autistic females carry a greater load of both rare *de novo* and common variants associated with autism than autistic males [6–8] and that recurrence likelihood is greater amongst male siblings than female siblings of autistic female probands [9].

Genetic studies have identified a large and heterogeneous set of genes associated with autism (https://gene.sfari.org/) [10]. Many of these autism-associated genes are highly expressed in neurons and have important roles in regulating synaptic transmission and plasticity [11,12]. Autism has also been associated with a number of developmental and environmental exposures including pregnancy complications [13] and *in utero* exposures [14]. Maternal infection during pregnancy has been shown to significantly increase the likelihood of autism in the offspring [15] and autistic individuals are more frequently observed to have immune dysregulation, including 36% greater odds of having an autoimmune condition [16]. Epigenetic mechanisms, including DNA methylation, are influenced by genetic variation, environmental exposures and developmental processes and play a role in regulating cell-type-specific gene expression. They therefore provide a potential molecular interface through which these factors contribute to autism.

Previous epigenome-wide association studies (EWASs) of DNA methylation variation associated with autism have primarily examined peripheral tissues, including blood and placenta [17–24], or profiled bulk post-mortem cortex [25–27]. The cellular heterogeneity of bulk brain tissue represents an important limitation since differences in cellular composition may obscure molecular variation occurring in specific cell types. This is particularly important as previous transcriptomic studies have consistently identified dysregulation to neuronal and microglial pathways [28–32]. Recent cell-type-resolved methylomic studies have begun to address this limitation, but have either only included one sex [33] or not investigated sex differences [34]. Sex stratification is particularly important given the marked sex difference in autism prevalence, and to ensure that findings from molecular studies are relevant to both males and females [35].

In this study, we analysed cell-type-and sex-specific DNA methylation variation in the human prefrontal cortex (PFC) associated with autism. Using fluorescence-activated nuclei sorting (FANS) [36], we isolated neuron-enriched (NeuN+), oligodendrocyte-enriched (SOX10+), microglia-enriched (IRF8+) and astrocyte-enriched (NeuN-/SOX10-/IRF8-) nuclei from 24 autistic (70.8% male) and 23 non-autistic (69.6% male) donors. Within each cell type, we performed EWAS analyses to identify methylomic differences associated with autism and with the interaction between autism diagnosis and sex. We identified pronounced cell-type- and sex-specific methylomic differences, with the strongest effects occurring at X-linked sites in microglia from autistic females. To our knowledge, this is the most comprehensive analysis to date of sex-specific DNA methylation variation across purified cortical cell populations in autism, highlighting a potential role for altered X chromosome regulation in female microglia.

## RESULTS

### Cell-type-resolved DNA methylation profiling of prefrontal cortex from autistic and non-autistic donors

Fluorescence-activated nuclei sorting was used to isolate purified populations of neuron-enriched (NeuN+), oligodendrocyte-enriched (SOX10+), microglia-enriched (IRF8+) and triple negative (TN) astrocyte-enriched (NeuN-/SOX10-/IRF8-) nuclei (**Figure S1**; **Table S1**) from prefrontal cortex (PFC) tissue from 47 individuals (n = 24 autistic donors, n = 23 non-autistic controls, age range = 6 – 91 years) (**Figure 1A-B**; **Figure S2**; **Table 1**). There was no difference in the age or sex distribution between groups (mean age [years]: autism = 31.4, control = 40.1, p = 0.120, *two-tailed T-test*; male-to-female ratio: autism = 2.43:1, control = 2.29:1). Subsequently, DNA methylation was quantified across the genome using the Illumina EPICv2 microarray (**Methods**) and, following stringent pre-processing, the final dataset included DNA methylation data for 857,146 DNA methylation sites (836,537 autosomal sites and 20,609 on the X chromosome) for each nuclei fraction. Principal component analysis (PCA) and hierarchical clustering of the 10,000 most variable sites across all purified nuclei populations showed complete separation of samples by cell type (**Figure 1C**; **Figure S3**), highlighting the distinct molecular profiles of the isolated nuclei populations. The purity of each cell type was confirmed experimentally using single-nucleus RNA sequencing from four donors, which demonstrated the expected enrichment of cell-type-specific marker gene expression within each nuclei fraction (**Figure S4**; **Methods**). Additionally, the cellular composition of each nuclei fraction was estimated computationally using a cortex-specific reference panel [37,38], which confirmed enrichment of the expected cell types in each nuclei fraction (**Figure S5**), with no difference in purity between diagnostic groups or sex (**Table S2**).

**Figure 1.**
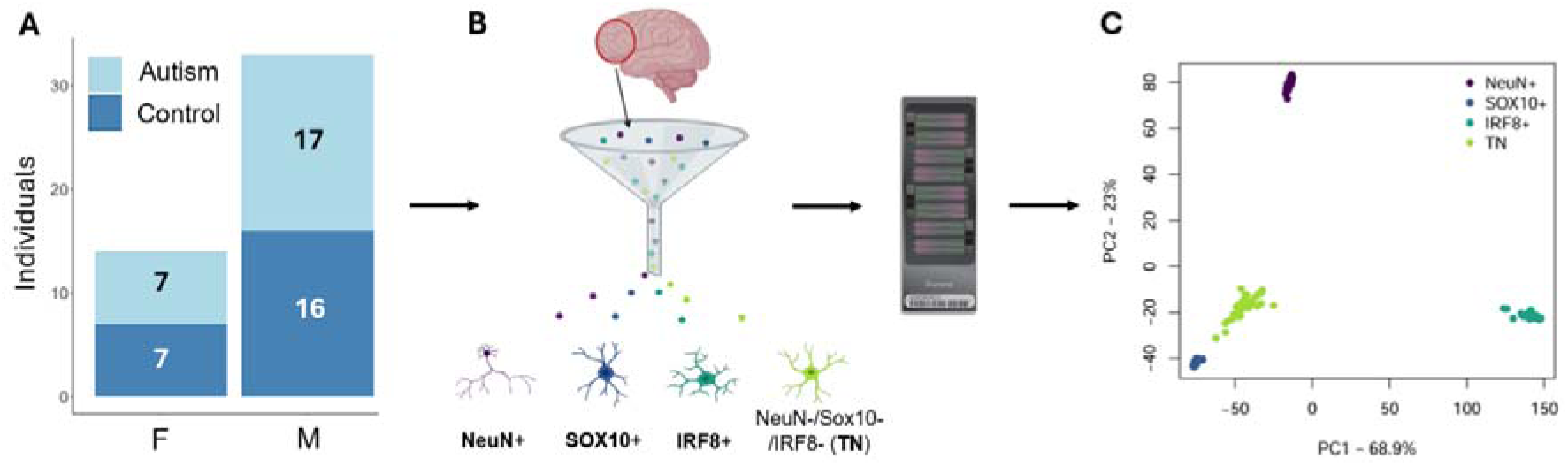
Generation of cell-type-resolved genome-wide DNA methylation data from the prefrontal cortex (PFC) of individuals with and without an autism diagnosis. **A)** Distribution of the 47 individuals included in this study stratified by sex and autism diagnosis. **B)** Schematic diagram showing the isolation of purified nuclei populations from PFC tissue and subsequent genome-wide DNA methylation profiling. Purified populations of NeuN+ (neuron-enriched), SOX10+ (oligodendrocyte-enriched), IRF8+ (microglial-enriched) and triple negative (TN) astrocyte-enriched (NeuN-/SOX10-/IRF8-) nuclei were isolated using fluorescence-activated nuclei sorting (FANS). DNA methylation profiles were generated for each cell fraction using the Illumina EPICv2 microarray. **C)** Principal component analysis (PCA) on the 10,000 most variable DNA methylation sites demonstrates clustering of samples by purified nuclei fraction, confirming the isolation of specific, non-overlapping cell type populations. The variance explained by the first two principal components is shown in the axis labels.

**Table 1.** Demographic overview of the study cohort.

|  |  |  |  |  |
| --- | --- | --- | --- | --- |
| Age range | 6 – 91 years |  |  |  |
| Sex | F |  | M |  |
|  | 14 |  | 33 |  |
| Age range | 13 – 91 years |  | 6 – 59 years |  |
| Autism diagnosis | Autism |  | Control |  |
|  | 24 |  | 23 |  |
| Age range | 6 – 91 years |  | 13 – 87 years |  |
| Sex | F | M | F | M |
|  | 7 | 17 | 7 | 16 |

### Autism-associated DNA methylation differences are cell-type-specific

To identify autism-associated differences in DNA methylation, we performed an epigenome-wide association study (EWAS) separately within each cell type by fitting linear regression models of DNA methylation as a function of diagnostic group (autism vs control), adjusting for sex and age (“autism EWAS”) (**Methods**). Although no DNA methylation sites passed an experiment-wide significance threshold (p < 9×10^-8^ [39]), 35 differentially methylated positions (DMPs), all located on the autosomes, were identified using a discovery threshold of p < 1×10^-5^. These DMPs were annotated to 29 unique genes (**Table S3**) and all were identified in only in a single cell type. All four cell types contributed at least one DMP, although more than half were identified in IRF8+ nuclei (number of autism DMPs: NeuN+ = 9, SOX10+ = 1, IRF8+ = 20, TN = 5) (**Figures 3A** and **3B**; **Figure S6**) suggesting that autism-associated DNA methylation differences are particularly enriched in microglia.

Autism DMPs mapped to genes with functional relevance to autism. In NeuN+ nuclei, DMPs included sites annotated to *TENM3* and *CABP7*, genes involved in neuronal migration, synapse formation and calcium signalling [40–42] (**Figure S7A**). The single oligodendrocyte-associated DMP mapped to *CRTC3*, a CREB-regulated transcriptional co-activator implicated in cellular energy metabolism [43,44] (**Figure S7B**). Astrocyte-associated autism DMPs include sites annotated to *CAMKK1* (**Figure S7D**). *CAMKK1* encodes a component of the CaMK4 signalling pathway, which has been associated with autism [45]. In contrast, many microglia-associated DMPs were annotated to genes with established roles in immune regulation and inflammatory signalling, including *GAPLINC* [46], *TNFSF10* [47], *TNRC18* [48,49], *LINC02068* [50,51], *FCGR3A* [52] and *TMEM163* [53,54] (**Figure S7C**). These findings are consistent with previous epigenetic and transcriptomic evidence implicating microglial and immune dysregulation in the cortex of autistic individuals [55–57].

Autism-associated effect sizes at the 35 DMPs showed limited concordance across nuclei fractions. For example, effect sizes at the 20 IRF8+ DMPs were only weakly correlated with those observed in NeuN+ (r = 0.125) and SOX10+ (r = 0.0980) nuclei and showed a modest but negative correlation with those in TN nuclei (r = −0.222). To formally assess cell-type-specificity, we re-analysed all 35 DMPs using a model incorporating all four cell types and an interaction term between autism diagnosis and cell type (**Methods**). Three DMPs demonstrated cell-type-specific differential methylation associated with autism. Two DNA methylation sites – cg20029967 (annotated to *SEPTIN11*, mean autism-associated DNA methylation difference = 3.26%, p = 9.06×10^-6^, **Figure 2A**) and cg13056990 (annotated to *AJAP1*, mean autism-associated DNA methylation difference = 3.82%, p = 5.92×10^-6^, **Figure 2B**) – showed neuron-specific differences in DNA methylation. *SEPTIN11* plays a role in GABAergic synaptic connectivity [58] and has previously been implicated in schizophrenia and bipolar disorder [59]. *AJAP1* is regulator of presynaptic inhibition and mutations in this gene have been associated with epilepsy, intellectual disability and neurodevelopmental disorders [60,61]. A third DMP (cg04410959 annotated to *TTC8*) exhibited a 13.5% autism-associated difference specific to the microglia-enriched nuclei (p = 2.68×10^-6^) (**Figure 2C**). *TTC8* encodes a component of the BBSome complex and pathogenic variants in the gene cause Bardet-Biedl syndrome (BBS), a ciliopathy with multisystem impacts including intellectual disability [62,63].

**Figure 2.**
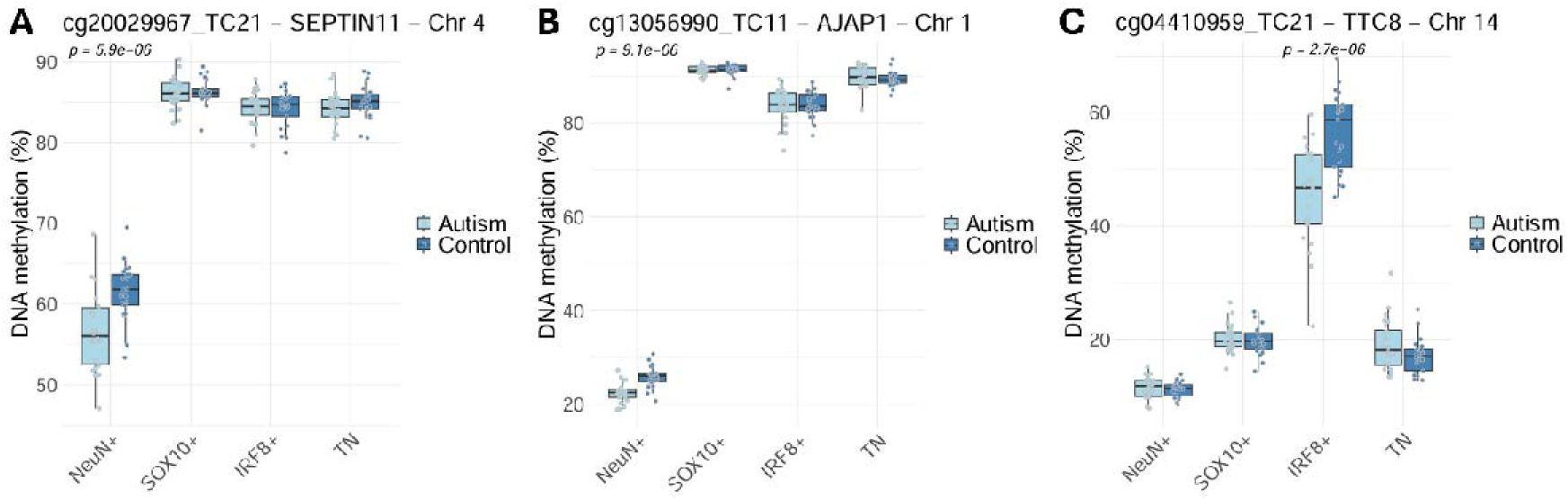
Cell-type-specific differences in DNA methylation associated with autism. **A)** NeuN+-specific difference in DNA methylation at cg20029967, characterised by hypomethylation in the NeuN+ nuclei of autistic individuals. cg20029967 is annotated to SEPTIN11 which is important for GABAergic synaptic connectivity [58]. **B)** cg13056990, annotated to AJAP1 – a regulator of presynaptic inhibition – demonstrates NeuN+-specific hypomethylation in autistic individuals. **C)** IRF8+-specific hypomethylation at cg04410959 in IRF8+ nuclei from autistic individuals. cg04410959 is annotated to TTC8, the protein of which it encodes forms a component of the BBSome complex. Linear regression p-values are shown for cell types in which the autism association reached nominal significance (p < 0.05).

Across all 857,146 sites tested, autism-associated effect sizes were generally correlated between the non-microglial fractions (**Figure S8**) but showed little concordance between IRF8+ nuclei and the other cell-types (correlation with IRF8+ autism effect sizes: NeuN+ (r = 0.0655); SOX10+ (r = 0.0830); TN (r = 0.0295)). This suggests that autism-associated methylomic variation in microglia is distinct from that observed in other cortical cell populations.

Finally, we compared cell-type-resolved autism effect sizes with those observed at 31 DMPs identified in our previous bulk prefrontal cortex study of idiopathic autism [27]. Among all four cell types, only IRF8+ nuclei showed a significant correlation with the bulk cortex effect sizes (correlation of autism-associated effect sizes identified in bulk cortex and IRF8+ nuclei = 0.41, linear regression p = 0.03) (**Figure S9**). This suggests that microglia account for a substantial proportion of the autism-associated DNA methylation differences previously detected in bulk cortex.

### Sex-by-autism interactions identify an X chromosome methylation signature in autism

To assess whether autism-associated DNA methylation differences differed between males and females, we fitted linear regression models that included a sex-by-diagnosis interaction term (**Methods**). At a discovery threshold of p < 1×10^-5^, 59 sites (annotated to 47 unique genes) demonstrated a significant interaction between sex and autism (number of sex-by-autism DMPs [of which X-linked]: NeuN+ = 11 [4], SOX10+ = 12 [9], IRF8+ = 28 [23], TN = 8 [2]) (**Figures 3A** and **3C**; **Figures S10** and **S11**; **Table S4**). Notably, IRF8+ nuclei were the only fraction to demonstrate experiment-wide significant interactions (p < 9×10^-8^), with 5 DMPs annotated to *MID1*, *PHF6*, *NRK*, *NONO* and *SAT1-DT*, all located on the X chromosome (**Figure 3B**). The autism-associated effects at these sites were substantially larger in females than males (mean absolute autism-associated DNA methylation difference [%] across the 5 DMPs [±SD]: females = 11.3 ± 1.43; males = 0.488 ± 0.644), suggesting a pronounced female-specific epigenetic signature within microglia that is absent in autistic males. The convergence of these effects on the X chromosome and in IRF8+ nuclei is also notable given our understanding of autism likelihood and the abundance of brain-expressed genes on the X chromosome, including many dosage-sensitive genes that are expressed from the Xi [64], and growing evidence implicating microglial dysfunction in autism [55–57].

**Figure 3.**
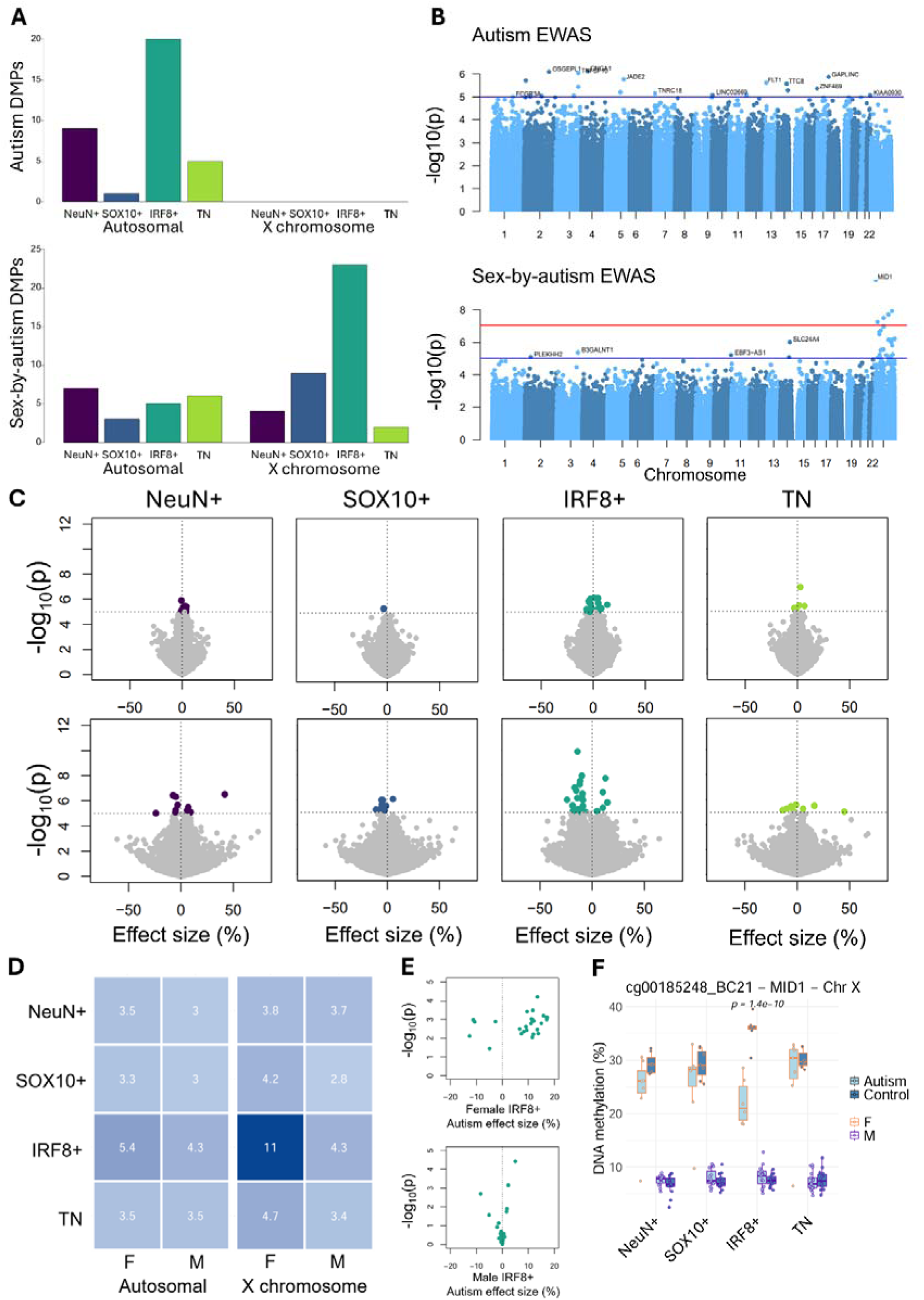
Autism-associated DNA methylation variation shows cell-type- and sex-specific differences. **A)** Differentially methylated positions (DMPs) identified in the autism and sex-by-autism EWAS analyses performed separately in each purified nuclei population, stratified by genomic location (autosomal or X chromosome). **B)** Manhattan plot showing the genomic distribution of -log10-transformed p-values identified in IRF8+ nuclei from the autism and sex-by-autism EWAS analyses. **C)** Volcano plots showing effect sizes and p-values from the autism EWAS (top) and sex-by-autism EWAS (bottom) for each tested DNA methylation site in each cell type. The horizontal dotted line indicates the discovery p-value threshold used to define a DMP (p = 1×10^-5^), with non-significant sites below this threshold coloured grey. **D)** For each set of sex-by-autism DMPs identified within a cell type (number of sex-by-autism DMPs: NeuN+ = 11, SOX10+ = 12, IRF8+ = 28, TN = 8), a heatmap showing the mean absolute autism effect sizes (mean autism-associated DNA methylation difference [%]) in males and females, stratified by genomic location (autosomal or X chromosome). Darker shades indicate greater magnitude of difference. **E)** At the 28 sex-by-autism DMPs identified in IRF8+ nuclei, volcano plots of autism effect sizes and p-values calculated in female and male IRF8+ nuclei separately. **F)** The top sex-by-autism DMP, cg00185248, annotated to MID1 on chromosome X, demonstrates a 13.5% mean difference in percentage DNA methylation between autistic and control females in IRF8+ nuclei.

The strongest interaction was observed at cg00185248, annotated to *MID1*, which was characterised by a larger autism-associated DNA methylation difference in females than males in IRF8+ nuclei (interaction p = 1.38×10^-10^, mean autism-associated DNA methylation difference = 13.5%, p = 6.14×10^-5^ [females]; −0.264%, p = 0.696 [males]; **Figure 3F**). *MID1* (*Midline-1*) encodes an E3 ubiquitin ligase required for typical neurodevelopment, including by mediating axonal growth and branching [65]. Pathogenic variants in *MID1* lead to Opitz BBB/G syndrome (OS), a congenital disorder affecting the development of midline structures during embryogenesis [66]. Owing to the buffering effect of X chromosome inactivation (XCI) in females, the severity of the OS phenotype demonstrates sexual dimorphism with males typically more severely affected than females [66]. The DNA methylation profiles observed at *MID1* (**Figure 3F**), and many of the other X-linked sex-by-autism DMPs (**Figure S10C**), resemble the canonical pattern expected for X-chromosome dosage regulation. Control females tend to exhibit intermediate DNA methylation levels (∼50%) consistent with the combined signal from an unmethylated active X chromosome (Xa) and a methylated inactive X chromosome (Xi). In contrast, autistic females show lower DNA methylation, potentially reflecting partial loss of methylation on the Xi. This observation is particularly intriguing given that a recent study of OS using iPSC-derived brain organoids demonstrated reactivation of the wildtype unmutated allele of *MID1* from the Xi during neural differentiation, directly modifying disease severity in a sex-dependent manner [67].

Many of the other sex-by-autism DMPs identified in IRF8+ nuclei (p < 1×10^-5^) were annotated to genes previously implicated in autism, including *EBF3* (SFARI Score 1S) [68,69], *TSPAN7* (SFARI Score 2) [70], *APOO* [71] and *WDR45* [72,73]). Also of interest is cg21966410, annotated to the androgen receptor gene, *AR*, which shows hypomethylation of autistic female IRF8+ nuclei (p = 1.66×10^-5^) (**Figure S12**). In NeuN+ nuclei, notable sex-by-autism DMPs included sites annotated to *CITED1*, a regulator of estrogen-dependent transcription [74] and *MAOA* (*Monoamine oxidase A*), a high-confidence autism-associated gene involved in neurotransmitter metabolism [75]. In SOX10+ nuclei two DMPs were annotated to *BCOR*, with both demonstrating hypomethylation in autistic females (mean autism-associated DNA methylation difference in females = 4.40%, p = 6.60×10^-4^ [cg06010380]; 11.0%, p = 3.79×10^-3^ [cg15039826]) and no difference in males (mean difference in males = −0.923%, p = 0.0775 [cg06010380]; −0.136%, p = 0.747 [cg15039826]). *BCOR* encodes the *BCL-6* transcriptional corepressor responsible for early embryonic gene expression regulation [76] and a paralog of *BCOR* (*BCORL1*) is associated with syndromic forms of autism [77]. Interestingly, one of these loci (cg15039826) showed comparable female-specific autism-associated methylation differences in NeuN+ (mean difference in females = 9.05%, p = 0.0133), SOX10+ (11.0%, p = 3.79×10^-3^) and TN (9.15%, p = 0.0336) nuclei, but not in female IRF8+ nuclei (0.185%, p = 0.969) (**Figure S10B**), suggesting that some female-specific epigenetic alterations may arise early during neural lineage specification before diverging between mature cell types.

We next formally tested the cell-type-specificity of the 59 sex-by-autism DMPs using a model incorporating all four nuclei populations simultaneously (**Methods**). 17 DMPs demonstrated significant cell-type-specific effects (**Table 2**), of which 14 were specific to IRF8+ nuclei. These included sites annotated to genes involved in epigenetic regulation (e.g. *OGT*) [78], axonal pruning (e.g. *XIAP*) [79] and immune signalling (e.g. *TSPAN7*) [80]. Together, these findings identify a predominantly X-linked, female-driven methylation signature that is concentrated in the microglia-enriched nuclei population.

**Table 2.** Cell-type-specific DMPs associated with an interaction between sex and autism diagnosis. Linear regression statistics for the 17 cell-type-specific sex-by-autism DMPs identified in NeuN+ (neuron-enriched), IRF8+ (microglia-enriched) and TN (astrocyte-enriched) nuclei populations. For each, the regression estimate and p-value are provided for the sex-by-autism interaction term, alongside the autism estimate and p-value calculated within females and males separately. The chromosome (CHR) and genomic position are provided, alongside the gene annotation for sites within 1500bp upstream of a gene.

| Cell type | Probe ID | CHR | Position | Gene | Sex-by-autism Estimate (%) | Sex-by-autism p-value | Female Autism Estimate (%) | Female Autism p-value | Male Autism Estimate (%) | Male Autism p-value |
| --- | --- | --- | --- | --- | --- | --- | --- | --- | --- | --- |
| IRF8+ | cg23949973_BC21 | X | 105821803 | NRK | 12.7 | 1.94E-08 | -11.2 | 1.04E-03 | 1.63 | 1.81E-02 |
| IRF8+ | cg21640139_BC21 | X | 71254819 | NONO | -11.6 | 3.11E-08 | 11.5 | 1.23E-03 | -0.108 | 7.00E-01 |
| IRF8+ | cg20308511_TC21 | X | 23782939 | SAT1-DT | -12.1 | 5.66E-08 | 11.0 | 1.62E-03 | -0.161 | 5.66E-01 |
| IRF8+ | cg18530240_BC21 | X | 71534447 | OGT | -16.9 | 1.05E-07 | 15.8 | 6.37E-04 | -0.481 | 6.63E-01 |
| IRF8+ | cg06609793_BC11 | X | 49080031 | WDR45 | -15.4 | 1.83E-07 | 15.0 | 1.57E-03 | -0.331 | 6.44E-01 |
| IRF8+ | cg10864286_BC21 | X | 35919832 | CFAP47 | 10.1 | 2.47E-07 | -10.7 | 1.36E-03 | 0.0148 | 9.43E-01 |
| IRF8+ | cg15170964_BC21 | X | 38561165 | TSPAN7 | -9.33 | 3.26E-07 | 9.04 | 2.38E-03 | -0.472 | 2.08E-01 |
| IRF8+ | cg26540943_TC11 | X | 155612635 | SPRY3 | -9.80 | 6.22E-07 | 9.36 | 3.71E-03 | -0.268 | 4.10E-01 |
| IRF8+ | cg06920429_TC21 | X | 132219474 | RAP2C | -10.5 | 7.26E-07 | 11.6 | 1.01E-03 | 0.363 | 3.78E-01 |
| IRF8+ | cg24195486_TC21 | 14 | 92322236 | SLC24A4 | -24.1 | 9.67E-07 | 17.0 | 7.77E-04 | -8.32 | 2.07E-03 |
| IRF8+ | cg27158037_BC21 | X | 69298946 |  | -18.8 | 2.85E-06 | 17.2 | 7.90E-04 | -1.32 | 4.78E-01 |
| TN | cg15215519_BC21 | 4 | 184293479 |  | -9.96 | 5.19E-06 | 7.25 | 1.93E-03 | -2.33 | 3.11E-02 |
| IRF8+ | cg24691854_TC11 | X | 140510997 |  | -13.8 | 5.61E-06 | 13.8 | 5.60E-03 | 0.579 | 2.77E-01 |
| NeuN+ | cg25429171_TC21 | 12 | 49449552 | SPATS2 | 5.82 | 5.63E-06 | -3.78 | 5.71E-04 | 2.07 | 6.59E-03 |
| IRF8+ | cg17033281_TC11 | X | 123860126 | XIAP | -11.9 | 7.17E-06 | 11.5 | 8.84E-03 | -0.364 | 2.65E-01 |
| IRF8+ | cg15673990_TC21 | 14 | 86069980 | LINC02328 | -18.5 | 8.05E-06 | 12.1 | 1.18E-03 | -5.13 | 2.69E-02 |
| NeuN+ | cg07607131_BC21 | 1 | 25026870 |  | -24.0 | 9.55E-06 | 13.8 | 3.93E-03 | -9.72 | 1.95E-03 |

### X-linked autism-associated differences are greatest in female microglia

We next estimated autism-associated effect sizes separately in males and females at each of the 59 sex-by-autism DMPs. Across all four nuclei fractions, autism effect sizes at X-linked DMPs were larger in females than in males, with the greatest difference observed in IRF8+ nuclei. This suggests that sex-by-autism DMPs in IRF8+ nuclei are predominantly driven by differences in females, with a mean difference in DNA methylation of 11% between autistic and control females at X chromosome DMPs, compared to a mean difference of 4.3% in males (**Figures 3D** and **3E**).

To determine whether this pattern extended beyond the DMPs identified in the interaction analysis, we performed an autism EWAS separately in males and females for each cell type (**Methods**). **Figure 4A** presents the mean autism effect size (mean autism-associated DNA methylation difference) by cell type and sex, stratified by genomic location (autosomal or X chromosome), providing an overview of global sex-specific patterns in autism-associated DNA methylation. Across all cell types, autism effect sizes were significantly larger in females than in males, regardless of genomic location (**Tables S5** and **S6**). The most pronounced sex difference was observed at X-linked sites in IRF8+ nuclei where the mean autism effect size was substantially greater in females than in males (difference in mean absolute female IRF8+ autism effect size and mean absolute male IRF8+ autism effect size at X chromosome sites [%] = 1.17, p < 1×10^-320^, *two-tailed T-test*) (**Figure 4B**). Overall, autism effect sizes were only weakly correlated between males and females, with correlations ranging from r = 0.0604 in TN to r = 0.0336 in SOX10+ at autosomal sites and r = 0.0205 in IRF8+ to r = 0.0326 in NeuN+ at X-linked sites (**Figure S13**).

**Figure 4.**
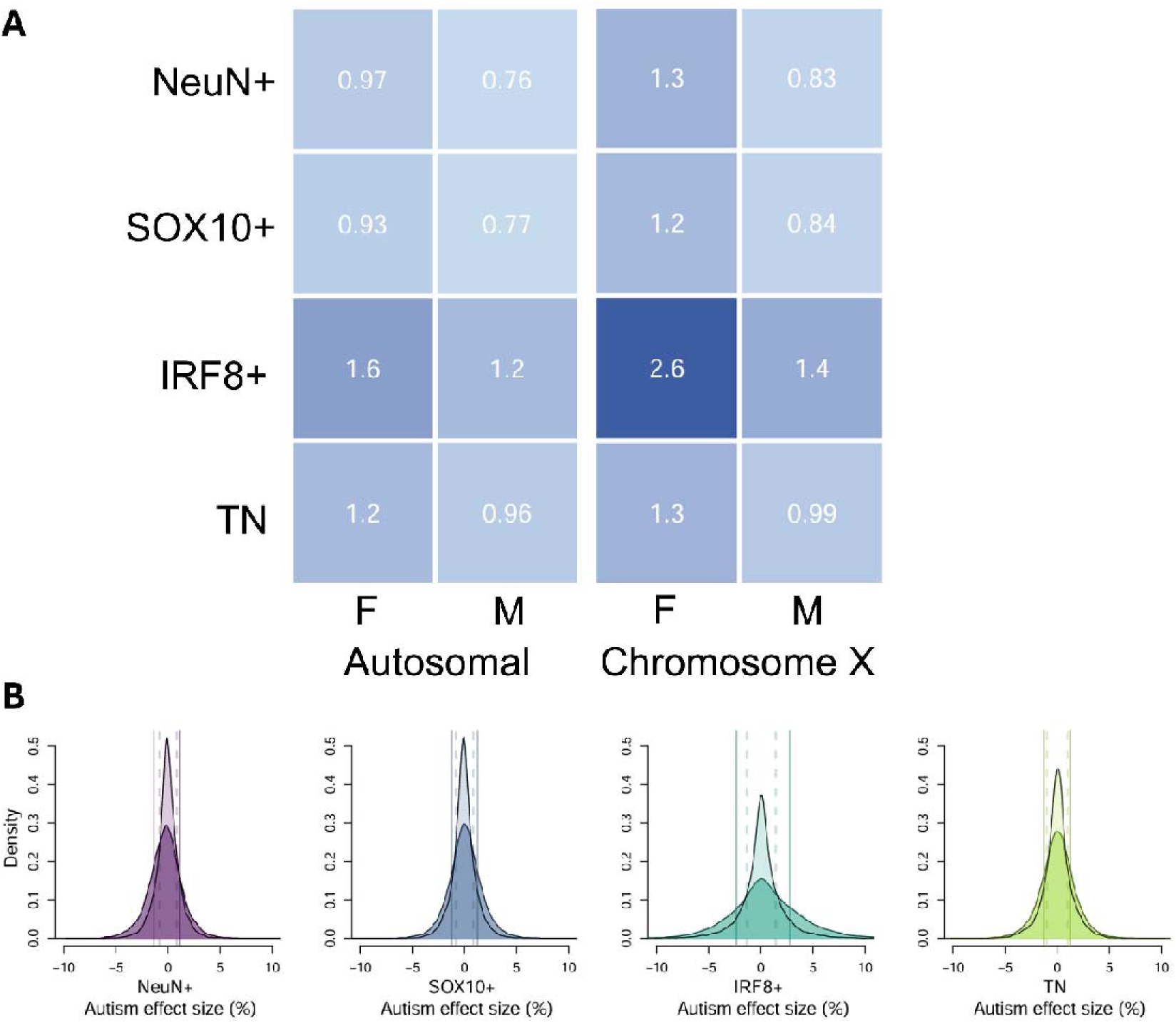
The largest autism-associated effect sizes are observed in female IRF8+ nuclei at sites on the X chromosome. **A)** Heatmap showing the mean absolute autism effect size (mean autism-associated DNA methylation difference [%]) in males and females, stratified by genomic location (autosomal or X chromosome). Darker shades indicate greater magnitude of difference. **B)** Autism effect sizes are significantly greater in females than in males on the X chromosome. Density plots showing the autism effect size in females (darker shades) and males (lighter shades) at sites on the X chromosome. Solid line = female mean; dotted line = male mean. Two-tailed T-test comparing female and male means is significantly different for all cell types, with IRF8+ nuclei having the largest difference (**Table S6**).

These results demonstrate widespread differences in the magnitude and direction of autism-associated methylomic variation between males and females. Autism effect sizes are consistently larger under three conditions and in an additive manner: in females, in IRF8+ nuclei, and at X chromosome sites. Hence, the largest effect sizes were observed in female microglia at X-linked sites (**Figure 4A**). This supports our previous observation of X-chromosome enrichment in the sex-by-autism EWAS and indicates that there is a broad X-chromosome-wide shift in autism-associated DNA methylation in female microglia. These results emphasise the importance of explicitly modelling sex-specific effects, rather than simply adjusting for sex, when studying sexually dimorphic phenotypes such as autism.

### Female-specific X-linked differences are consistent with altered X chromosome inactivation

Given the enrichment of female-specific autism-associated DNA methylation differences on the X chromosome, we next explored whether these effects differed according to XCI status. Using a consensus set of 107 genes that escape XCI and remain actively transcribed from the inactive X chromosome (Xi) [81], X-linked DNA methylation sites were classified according to whether they were annotated to genes that escape XCI or to genes normally subject to XCI (**Methods**). Across all cell types, except TN nuclei, mean absolute autism effect size in females were significantly greater at sites annotated to genes subject to XCI than at sites annotated to escape genes (p < 0.05, *two-tailed T-test*; **Figure 5A**; **Table S7**). The largest difference was observed in IRF8+ nuclei where the mean absolute autism-associated effect size was 2.61% at non-escape genes compared with 2.23% at escape genes (p = 4.15×10^-13^, *two-tailed T-test*). Although this difference is modest at individual loci, this shift in DNA methylation level over thousands of sites across the X chromosome is consistent with a widespread perturbation to gene dosage regulation at genes normally subject to XCI, particularly in microglia. Furthermore, many X-linked sex-by-autism DMPs in IRF8+ nuclei showed lower DNA methylation in autistic females than in non-autistic females (**Figure 5B**). Similar loss of Xi-associated DNA methylation has been reported in other sex-biased disorders such as lupus [82], systemic sclerosis [83] and breast cancer [84].

**Figure 5.**
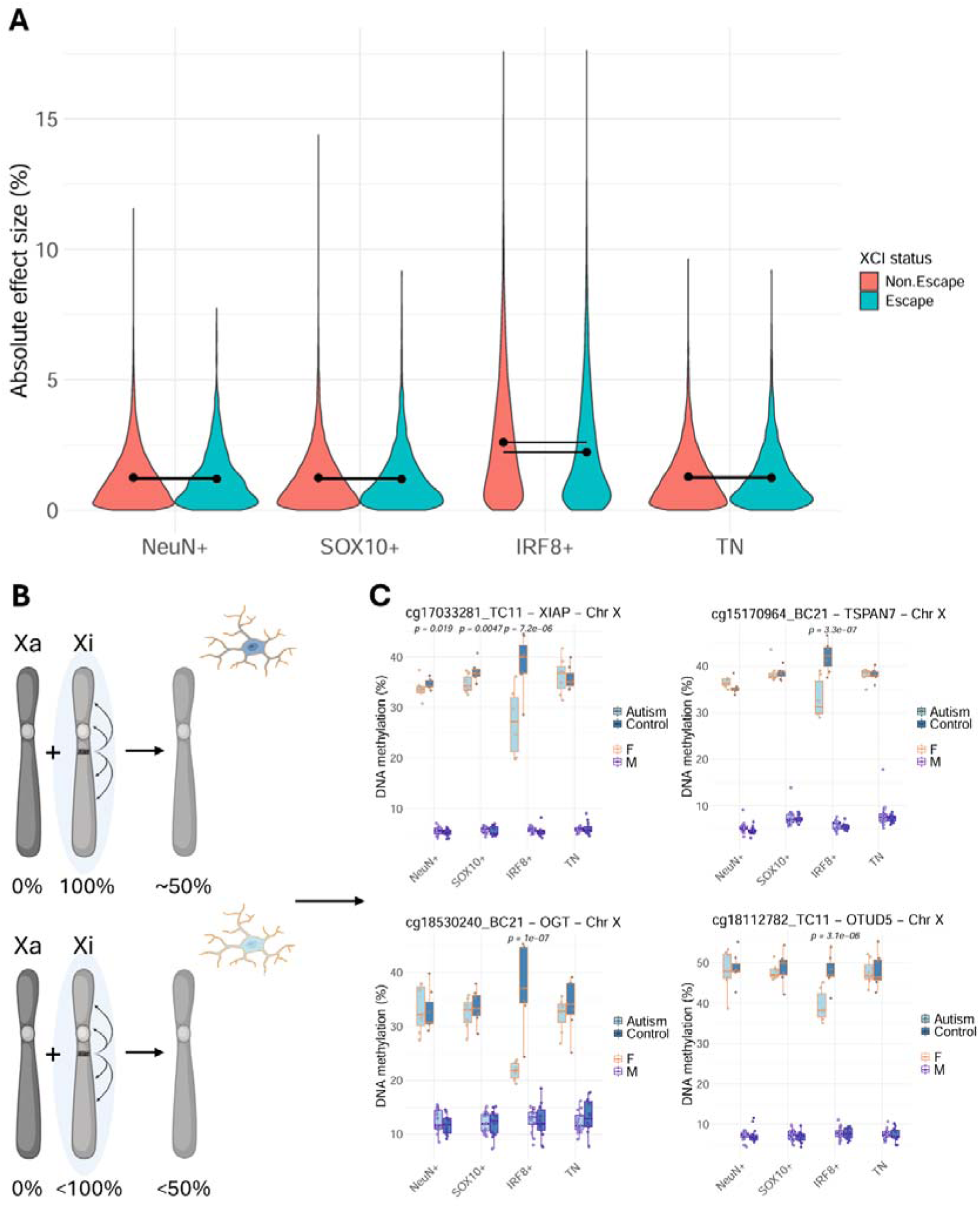
Autism-associated methylomic variation in female microglia may reflect loss of DNA methylation from the inactivate X chromosome. **A)** Distribution of absolute autism effect sizes (mean autism-associated DNA methylation difference [%]) in females for X-linked sites, stratified by the XCI status of their annotated genes. Solid circles denote mean absolute effect sizes and are connected within a cell type by solid lines. **B)** Illustration of proposed loss of DNA methylation from the inactive X chromosome (Xi) in autistic (light blue) female (orange outline) microglia compared to control (dark blue) females. **C)** Observations of lower DNA methylation in microglia from autistic females compared to control females in four representative examples of the six X-linked genes found to have higher expression in autism frontal cortex [32].

To investigate the potential functional relevance of these female-specific DNA methylation differences, we investigated autism-associated gene expression changes for genes annotated to X-linked sex-by-autism DMPs identified in IRF8+ nuclei. Nineteen of the 23 DMPs were annotated to a gene and could be evaluated using the ASD Single-Cell Gene Expression Portal (http://solo.bmap.ucla.edu/asdscgene/) [32]. Six genes (*TSPAN7*, *XIAP*, *OTUD5*, *OGT*, *NONO* and *ELF4*) showed significantly higher expression in autism (FDR < 0.05) in one or more microglia cell sub-types (MG1-homeostatic and MG2-reactive) (**Figure 5C**). Three genes (*MID1*, *MAGIX and SPRY3*) showed significantly lower expression (**Table S8**). These findings suggest that several genes linked to female-specific DNA methylation differences in microglia are also differentially expressed in the autistic cortex.

### Autism-associated co-methylation networks implicate neurodevelopmental and immune pathways

To identify co-ordinated DNA methylation changes associated with autism we performed weighted gene correlation network analysis (WGCNA) [85] separately within each nuclei fraction (**Methods**). Between 18 and 33 co-methylation modules were identified per fraction (**Table S9**) with the sites annotated to each module reported in **Table S10**. The first principal component of each module (the “module eigengene”) was tested for association with autism diagnosis and, in a separate model, the interaction between sex and diagnosis. Eight modules were nominally associated with autism diagnosis (number of significant modules: NeuN+ n = 2, SOX10+ n = 1, IRF8+ n = 1, TN n = 4; p < 0.05) with an additional module in IRF8+ nuclei associated with the sex-by-autism interaction (p < 0.05; **Figure 6**; **Table 3**).

**Figure 6.**
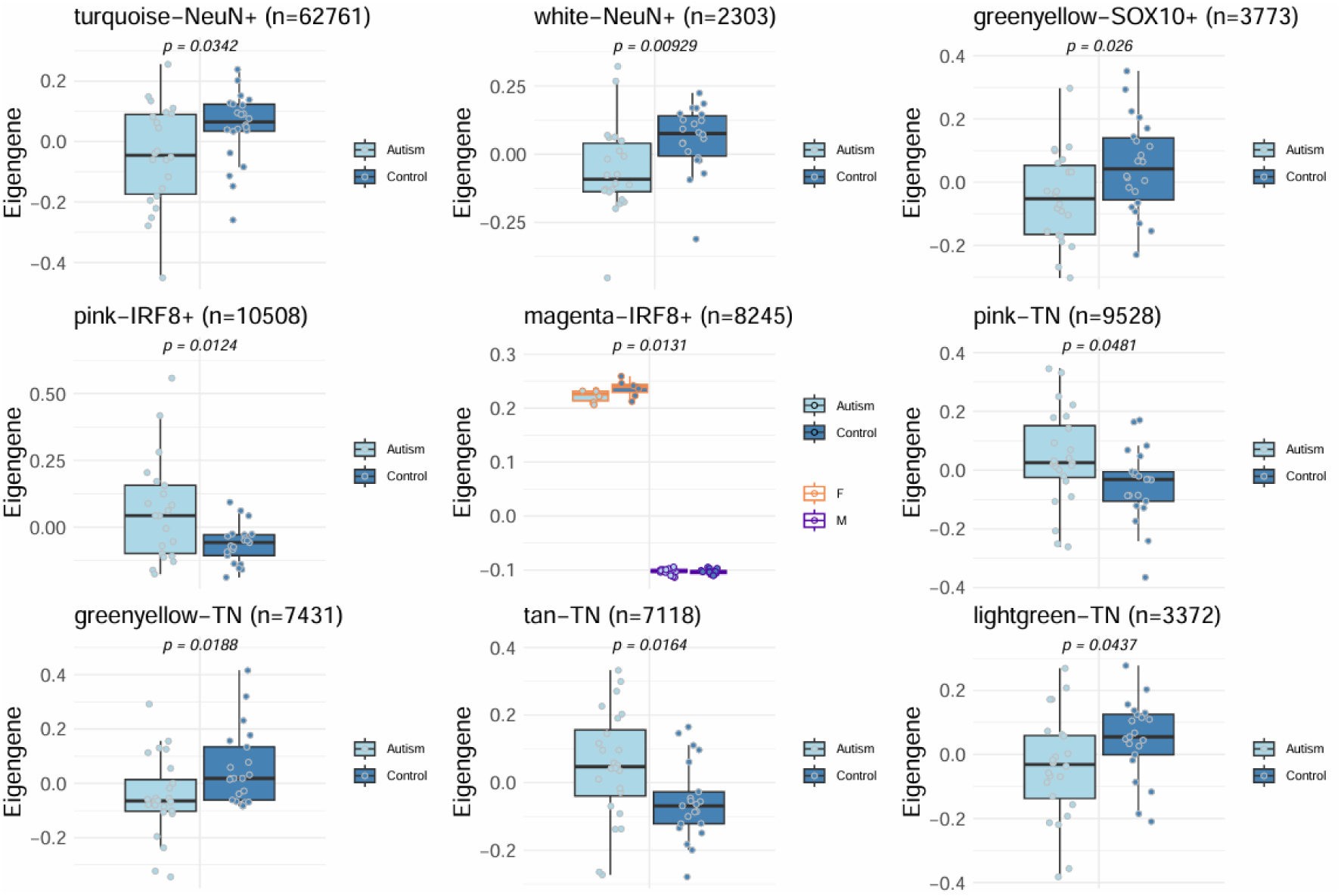
Co-methylation network modules associated with autism. Nine co-methylation network modules with a significant association between the module eigengene and autism diagnosis or the interaction between autism and sex (p < 0.05). The magenta-IRF8+ module was the sole module significantly associated with a sex-by-autism interaction. The total number of sites within the module is indicated in the plot title.

**Table 3.**
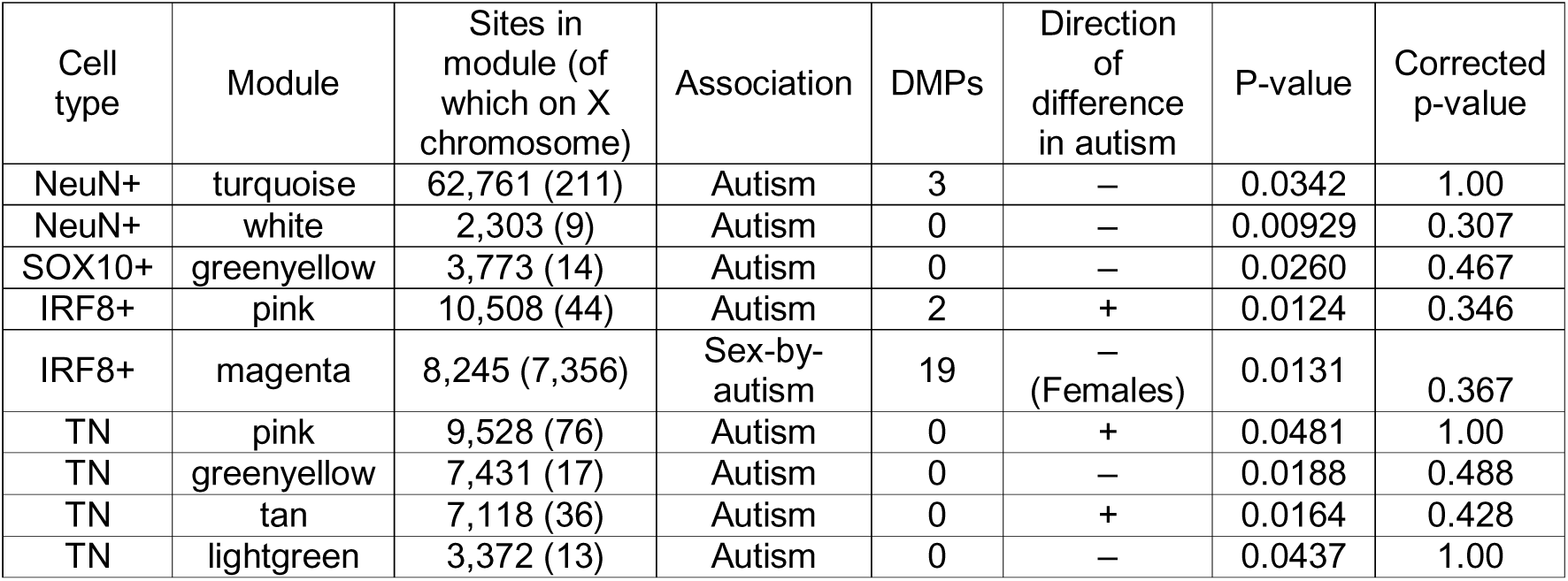
Co-methylation network modules associated with autism. WGCNA modules with a significant (p < 0.05) association between the first principal component of the module and autism diagnosis or a sex-by-autism interaction. The direction of effect (hypomethylated (‘–’) or hypermethylated (‘+’) in nuclei from autistic individuals) is shown alongside raw and adjusted p-values (corrected for the number of modules identified in each cell type). The number of sites in each module that overlap differentially methylated positions (DMPs) identified in the associated EWAS (autism EWAS or sex-by-autism EWAS) are shown.

For each associated module, module membership was calculated for every DNA methylation site to identify highly connected hub sites. Module membership was significantly correlated with autism-associated EWAS effect sizes (**Figure S14**), indicating that the strongest autism-associated loci tended to be more highly connected within the associated co-methylation networks (**Figure S15**). The 20 most highly connected hub sites for each module included sites annotated to several genes previously implicated in autism and/or intellectual disability (e.g. pink-IRF8+: *WNT2B* [86] and *GNAI2* [87]; magenta-IRF8+: *PQBP1* [88], *MAP7D3* [89–91], *NONO* [92,93] and *MAMLD1* [94]; pink-TN: *ZC3H12C* [95] and *CARS1* [96]). Hub sites were also annotated to several genes included in the Simons Foundation Autism Research Initiative (SFARI) Gene database of high-confidence autism-associated genes (https://gene.sfari.org/) (turquoise-NeuN+: *SHANK2* [Score 1]; magenta-IRF8+: *MSL3* [Score 1S], *CASK* [Score 1], *CUL4B* [Score 3S]; tan-TN: *PLXNB1* [Score 2], *NTRK2* [Score S], *PUF60* [Score 3S]; pink-TN: *CDC42BPB* [Score 2]; greenyellow-TN: *RPH3A* [Score 3], *KCNS3* [Score 2]).

The sole module (magenta) associated with the sex-by-autism interaction was identified in IRF8+ nuclei. Notably, 89.2% of the DNA methylation sites within this module mapped to the X chromosome, including 19 of the 23 X-linked sex-by-autism DMPs identified in IFR8+ nuclei. Consistent with our observation across the entire X chromosome (**Figure 4B**), across all 8,245 sites comprising the module, autism-associated DNA methylation differences were significantly greater in females than males (difference in mean autism-associated effect size [%] = 1.56, p < 1×10^-320^, *two-tailed T-test*) (**Figure S16**). This network-based analysis therefore provides additional support for co-ordinated sex-dependent methylomic variation across the X chromosome in microglia-enriched nuclei.

Gene Ontology (GO) analysis identified enrichment of associated modules for biological processes relevant to autism (**Figure S17**; **Table S11**). The turquoise-NeuN+ module, which showed significantly lower DNA methylation in autism, was enriched for neurodevelopmental pathways including “neurogenesis” (OR = 1.04, p = 1.07×10^-7^) and “nervous system development” (OR = 1.04, p = 3.93×10^-8^). This was the largest of the nine associated modules, containing over 60,000 sites. The pink-IRF8+ module, which showed significantly higher DNA methylation in autistic individuals, was enriched for immune-associated pathways including “immune response” (OR = 1.32, p = 8.99×10^-44^), “inflammatory response” (OR = 1.29, p = 5.60×10^-25^) and “response to cytokine” (OR = 1.24, p = 2.09×10^-17^) (**Figure S18**). In contrast, the predominantly X-linked magenta-IRF8+ module was enriched for synaptic pathways including “synapse organization” (OR = 1.07, p = 5.60×10^-6^) and “glutamatergic synapse” (OR = 1.09, p = 1.96×10^-6^). Given the abundance of brain-expressed genes on the X chromosome and the predominantly X-linked composition of this module, these enrichments should be interpreted cautiously. Nevertheless, the network analyses converge with the EWAS results in identifying neuronal, immune-related and female-specific X-linked methylation signatures associated with autism.

## DISCUSSION

This study provides a cell-type-resolved analysis of sex-specific DNA methylation variation associated with autism in the human prefrontal cortex, identifying female microglia as a major site of autism-associated epigenetic dysregulation. Across complementary locus-level, sex-stratified and network-based analyses, we observed convergent evidence for widespread X-linked DNA methylation differences in female microglia that are consistent with altered regulation of X chromosome inactivation.

Previous transcriptomic studies of autism have consistently identified dysregulation of neuronal and microglial pathways, characterised by a down-regulation of synapse-related transcripts and increased expression of microglial-mediated immune signalling pathways [28–32]. Our findings provide complementary epigenetic evidence supporting this neuroimmune signature. Neuron-specific autism-associated DMPs were annotated to genes involved in synaptic connectivity and regulation of presynaptic inhibition, which are highly relevant in the context of altered synaptic processes in autism [97], whereas microglial nuclei, which represented the largest group of autism-associated DMPs, were annotated to genes involved in immune function and inflammatory pathways. As specialised macrophages of the brain, microglia are essential for regulating neurodevelopmental processes such as synaptic pruning and neurogenesis [98–100] and their over-activation leading to neuroinflammation and impaired synaptic function are prominent features of autism [56,101]. Importantly, these observations were independently reinforced by our co-methylation network analyses. Of particular note is the enrichment of immune activation pathways associated with sites annotated to the pink-IRF8+ module, which exhibits significant hypermethylation of microglia from autistic donors. These findings are consistent with previous reports of autism-associated DNA methylation differences in microglia [33,34,102] and strengthen the growing evidence implicating immune dysregulation in autism [56,101].

The cellular heterogeneity of bulk tissue limits both the sensitivity to detect phenotype-associated molecular variation and the ability to identify the cell types in which such variation occurs. By profiling purified neural nuclei populations, we show that autism-associated effect sizes are highly cell-type-specific with weak concordance across cell types, particularly for microglia. This approach enabled the identification of neuronal- and microglial-specific differential methylation associated with autism and suggests that microglia are a major contributor to the autism-associated differences previously observed in bulk prefrontal cortex [27]. This same principle extends to the analysis of sexually dimorphic phenotypes, such as autism, where the use of combined- or single-sex cohorts may obscure biologically important sex-dependent molecular differences [103]. By explicitly modelling the interaction between sex and autism diagnosis, we identified a striking female-specific epigenetic signature that would have been otherwise missed. Together, these findings highlight the importance of considering both cellular composition and sex in molecular studies of autism.

Our results contribute to a growing body of epigenomic and transcriptomic evidence for female-specific molecular variation across neurodevelopmental and neuropsychiatric conditions, including autism [20,21], schizophrenia [104] and major depressive disorder [105]. Most sex-by-autism DMPs identified in microglia were located on the X chromosome with several annotated to genes previously associated with autism. At many of these sites, autistic females showed lower DNA methylation than non-autistic females in a pattern consistent with a loss of DNA methylation from the inactive X chromosome (Xi). Interrogation of results from a single-cell gene expression study of autism [32] confirmed that nine of the genes annotated to X-linked microglial sex-by-autism DMPs were significantly differentially expressed (six up-regulated, three down-regulated) in microglia from autistic donors. This overlap supports the potential functional relevance of the identified loci and raises the possibility that some microglial expression differences reported in combined-sex studies are influenced disproportionately by changes in autistic females. However, because the methylation and expression data were generated in different cohorts and the transcriptomic results were not stratified by sex, matched sex-stratified multi-omic studies will be required to determine whether these DNA methylation differences directly influence gene expression (Wamsley et al., 2024).

Across the X chromosome, autism-associated methylomic differences were significantly larger in females than males, particularly in microglia, and were enriched at genes normally subject to XCI compared to genes that escape XCI. Co-methylation network analysis reinforced the EWAS findings by identifying a predominantly X-linked microglial module associated with the sex-by-autism interaction. This module contained 19 of the 23 X-linked sex-by-autism DMPs and showed autism-associated hypomethylation in female microglia. The convergence of these locus-level, chromosome-wide, transcriptomic and network-based analyses provides strong evidence that female-specific autism-associated epigenetic variation is concentrated on the X chromosome and is consistent with XCI instability involving loss of Xi methylation. Although direct allele-specific measurements of Xi status are required to further test this hypothesis, our findings identify altered X chromosome dosage regulation in microglia as a plausible mechanism contributing to sex-specific differences in autism.

Among the genes annotated to female-specific autism-associated DMPs in microglia was *OGT*, which was also differentially expressed in microglia in the independent combined-sex transcriptomic dataset. *OGT* (O-linked N-acetylglucosamine transferase) encodes the sole enzyme that catalyses the addition of O-GlcNAc to nuclear and cytoplasmic proteins. It is a critical regulator of gene expression, notably including X chromosome inactivation via its interaction with the polycomb repressive complex 2 (*PRC2*) [106]. *PRC2* is required to maintain stability of XCI gene silencing marks in extra-embryonic tissue [107] and human microglia are understood to be derived from extra-embryonic erythromyeloid progenitors in the yolk sac [108]. The methylomic changes observed in *OGT* in this study could signal a dysregulation of XCI epigenetic maintenance in autistic females. This could potentially impact the dosage regulation of many X-linked genes and explain the substantial X-chromosome-wide differential methylation observed in autistic female microglia. Sex differences in *OGT* expression have also been found to mediate sex differences in placental vulnerability [109], highlighting the importance of OGT in explaining sex-specific vulnerability in early development.

Our results extend recent whole genome bisulfite sequencing (WGBS) studies of umbilical cord blood [20] and neonatal blood [21] that identified larger and more abundant autism-associated differentially methylated regions in females and an enrichment for regions annotated to the X chromosome. These findings have been interpreted within the framework of the female protective effect in autism whereby females require a greater molecular burden to surpass the diagnostic threshold [6,8]. The concentration of female autism-associated methylomic differences on the X chromosome suggests that there may exist an altered state of XCI in autistic females [20,21,110]. Our results extend these observations by demonstrating that this female-specific epigenetic signature is present in the postnatal prefrontal cortex and is particularly pronounced in microglia. Whether these methylomic changes are a cause, consequence or compensatory response to autism is yet to be determined but our results identify female microglia as a key cellular context in which to investigate the mechanisms underlying sex-specific vulnerability in neurodevelopment. This study has several limitations. First, although in line with previous cortical DNA methylation studies of autism [26,27,33,34,111,112], our sample size remains modest, particularly for analyses of females, reflecting the challenges in obtaining tissue from female autistic donors. We deliberately enriched our cohort for female donors resulting in a male-to-female ratio of approximately 2:1, compared with the estimated population sex ratio of 3:1 [4]. It is notable that all five experiment-wide significant sex-by-autism DMPs were detected in microglia, despite the limited sample size. Second, while the Illumina EPICv2 array enables precise quantification of DNA methylation at single-base resolution and targets substantially more DNA methylation sites than those used in previous studies of autism (27K, 450K and EPIC arrays), it covers only a small proportion of DNA methylation sites in the human genome. As sequencing costs decline, future research should utilise sequencing-based technologies to comprehensively profile DNA methylation differences in autism. Third, our study could not distinguish between DNA methylation and its oxidised form, DNA hydroxymethylation, due to limitations of sodium bisulfite-based approaches [113]. This distinction is potentially important as DNA hydroxymethylation is abundant in the central nervous system and known to impact gene expression [114], although levels are minimal in microglia and thus unlikely to account for the principal findings in this cell type. Future studies should aim to differentiate these modifications using techniques such as nanopore sequencing [115]. Fourth, while our nuclei sorting strategy substantially improves cellular resolution, the nuclei populations analysed here, particularly the neuronal and triple negative populations, will contain a heterogeneous mix of cell sub-types. Future studies should prioritise single cell approaches to investigate autism-associated methylomic variation, such as those conducted by Eyring *et al.* [33]. Fifth, we were unable to generate gene expression data from these samples, preventing direct conclusions about the transcriptional impact of the observed DNA methylation changes. However, by integrating results from a single-cell transcriptomic study [32] we were able to interpret sex-by-autism DMPs in the context of known autism-associated gene expression changes. Finally, the cross-sectional analysis of post-mortem tissue cannot establish when DNA methylation changes arise or whether they represent causal, downstream, or compensatory processes.

In summary, this study provides the first comprehensive cell-type-resolved characterisation of sex-specific DNA methylation differences associated with autism in the human prefrontal cortex. We identify female microglia as a major focus of autism-associated epigenetic variation with widespread X-linked DNA methylation differences that are consistent with altered regulation of X chromosome inactivation in the autistic female brain. These results further our understanding of methylomic variation in the human prefrontal cortex and emphasise the importance of cell-type-and sex-specific analyses in studies of autism.

## METHODS

### Isolation of cell-type-specific nuclei populations from post-mortem cortex tissue

Post-mortem prefrontal cortex (PFC) tissue was provided from the SFARI Autism BrainNet resource (https://www.sfari.org/resource/autism-brainnet/), a collaborative network of academic sites that collects, stores and distributes brain tissue for autism research. Subjects were approached in life for written consent for brain banking, and all tissue donations were collected and stored following legal and ethical guidelines. All samples were dissected by trained neuropathologists, snap-frozen and stored at −80°C before being transferred to the University of Exeter under a Material Transfer Agreement. An overview of the samples used in this study is provided in **Table 1**.

### Fluorescence-activated nuclei sorting of specific nuclei populations from the prefrontal cortex

∼500mg PFC tissue was processed using fluorescence-activated nuclei sorting (FANS) [36]. A full protocol detailing each step of our nuclei purification protocol is provided on protocols.io at https://dx.doi.org/10.17504/protocols.io.36wgq4965vk5/v5. Briefly, following tissue homogenization and nuclei purification using sucrose gradient centrifugation, we used a FACS Aria III cell sorter (BD Biosciences) to simultaneously collect populations of NeuN+ (neuronal-enriched) (Millipore, Cat No: MAB377X, dilution: 1:1000), SOX10+ (oligodendrocyte-enriched) (R&D systems, Cat No: NL2864R, dilution: 1:20) and IRF8+ (microglia-enriched) (Invitrogen, Cat No: 17-9852-82, dilution: 1:150) immunolabeled populations from bulk DLPFC tissue prior to genomic profiling, with the triple-negative fraction (astrocyte- and other glia-enriched) also being collected from each tissue sample. Nuclei suspensions were assessed for the presence of debris by adjusting the gating strategy before proceeding with nuclei capture. For each sorted population, ∼50,000 nuclei were collected for DNA methylation profiling [36,116]. Details of the gating strategies we implemented are shown in **Figure S1**.

### DNA methylation profiling, processing and quality control

Approximately 50,000 nuclei for each cell-type-resolved sample were processed using the Zymo EZ-96 DNA Methylation-Direct Kit (Cambridge Bioscience, UK) according to the manufacturer’s standard protocol. All samples were then processed using the Illumina Infinium MethylationEPIC v2.0 array (Illumina Inc, CA, USA) according to the manufacturer’s instructions, with minor amendments and quantified using an Illumina iScan System (Illumina, CA, USA). Individuals were randomised and sorted fractions from the same individual and FANS gating run were processed on the same BeadChip, where within a BeadChip the location of each fraction was randomised. DNA methylation data was loaded into R (version 4.2.1) from IDAT files using the package *bigmelon* R package [117] and processed through a bespoke quality control pipeline developed for cell-specific DNA methylation data (https://github.com/ejh243/BrainFANS) [118]. Samples with methylated signal intensity < 500, bisulfite conversion efficiency < 80%, percentage missing data > 2%, high normalisation violence (> 0.1) or > 1% of probes with detection p-value > 0.05 were excluded. Sample genetic relatedness was confirmed using the 65 ‘rs’ SNP probes on the array. Samples with correlation > 80% to another unrelated sample were excluded and samples with < 80% correlation to another related sample (i.e. a different cell type from the same individual) were excluded. Correct reported sex was confirmed using intensities of probes on the X and Y chromosomes. For each cell type, the mean and standard deviation were calculated for the first two principal components. Samples > 3 standard deviations from the mean were excluded. Probes with a beadcount < 3 and probes where > 1% of samples had a detection p-value of > 0.05 were excluded. Data were normalised within each cell type using the *adjustedDasen* function from the *wateRmelon* R package [119].

DNA methylation sites were filtered to be biologically and technically relevant. First, DNA methylation sites annotated to the Y chromosome were excluded due to the absence of a Y chromosome in females. Second, array probes overlapping single nucleotide polymorphisms (SNPs) [120,121], as well as autosomal probes that were previously found to have high potential to cross-hybridise to off-target sites on the X and Y chromosomes [122], were excluded. Third, replicate EPICv2 array probes were investigated using the manifest file developed by Peters et al. [122]. The ‘Rep results by NAME’ column was used to identify the optimal replicate probe to keep, such as those labelled ‘Superior probe’ or ‘Best sensitivity’. If no clear superior probe was present (for example, when all replicate probes were classified as ‘Insufficient evidence’), the first probe in the replicate probe set was kept. The final dataset contained 857,146 DNA methylation sites for subsequent analysis. Sites were annotated to genes using the *ChIPseeker* R package [123] (GencodeV44, hg38). Where sites overlap multiple transcripts, annotations were prioritised in the order as follows: promoter region (defined as within 1500bp of the transcription start site), 5’ untranslated region (5’UTR), 3’UTR, exon, intron, downstream, intergenic.

After quality control and normalisation, the dataset contained DNA methylation data at 857,146 sites for PFC tissue from 47 post-mortem donors (24 autistic [7 female, 17 male], 23 control [7 female, 16 male]) each with up to four profiled cell types (44 “NeuN+” [NeuN+], 44 “SOX10+” [NeuN-/SOX10+], 42 “IRF8+” [NeuN-/SOX10-/IRF8+], 45 “TN” [NeuN-/SOX10-/IRF8-]) (**Table S1**). The cellular composition of purified nuclei fractions was estimated for each sample using cortex-specific reference panels provided through the *CETYGO* R package [37] (reference panel 2 with ANOVA method). Sorting efficiency was defined as the predicted proportion of the labelled cell type. A two-tailed Student’s T-test was used to assess differences in sorting efficiency between diagnostic groups separately for each cell type.

### Single-nucleus RNA-sequencing

To further validate the purity of the sorted nuclei populations, we performed single-nucleus RNA sequencing (snRNA-seq) on PFC nuclei isolated from four donors. Minor modifications were made to the FANS protocol to optimise RNA integrity, including the addition of Ribolock RNase inhibitor (Thermo Scientific, EO0382; 0.2 U/ml) to both the lysis and staining buffers. For each nuclei fraction, 100,000 sorted nuclei were fixed using the Parse Evercode Low-Input Nuclei Fixation Kit (v3), with approximately 6,250 nuclei targeted for recovery per sample. snRNA-seq libraries were prepared using the Evercode WT Kit (v3) and library quality assessed using the Agilent D5000 High Sensitivity ScreenTape assay. Libraries were pooled and sequenced on an Illumina NovaSeq 6000 platform (100 bp paired-end reads) to a target depth of ∼20,000 reads per nucleus. Sequencing reads were aligned to the GRCh38 reference genome and filtered count matrices generated using Trailmaker (v1.5.1, Parse Biosciences), excluding barcodes containing fewer than 10 transcripts or failing the barcode rank plot threshold. Putative doublets were identified and removed using scDblFinder [124].

Downstream analyses were performed in Seurat (v5.2.1) [125]. Nuclei with at least 200 detected transcripts and fewer than 10% mitochondrial reads were retained, alongside genes detected in at least five nuclei. Data from each sample were log-normalised and scaled, with the top 4,000 variable features selected for dimensionality reduction by principal component analysis (PCA). To account for inter-sample technical variation, datasets were integrated using the *Harmony* [126] prior to graph-based clustering using the Leiden community detection algorithm. Uniform Manifold Approximation and Projection (UMAP) embeddings were generated for the visualisation of cellular structure. Following a second round of filtering excluding nuclei with fewer than 400 detected genes, clusters were manually annotated as major neural cell populations based on established marker gene expression profiles.

### Identifying differentially methylated positions associated with autism

To identify loci that demonstrate significant differences in mean DNA methylation levels between individuals with and without an autism diagnosis (*Group*), a linear regression model was fitted to each DNA methylation site, *i,* with covariates for *Sex* and *Age* (years). Analyses were performed separately for each cell type (“within-cell-type EWAS”) using the following model:

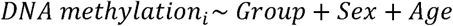

To identify differences associated with a sex-by-autism interaction, the model was expanded to include an interaction term between sex and diagnostic group (*Sex*Group*):

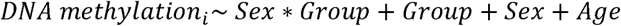

To identify autism-associated differences separately within each sex, the within-cell-type autism EWAS was performed separately in males and females using the following model:

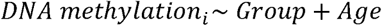

Differences in mean DNA methylation level at a given site were considered experiment-wide significant if the Student’s t-distribution p-value for the *Group* or *Sex*Group* covariate was < 9×10^-8^. This is considered a suitable threshold to account for multiple testing in epigenome-wide studies on the EPIC array in large case-control analyses [39]. However, due to the limited power in this study to identify genome-wide significant differences within each cell type, differentially methylated positions (DMPs) were defined as sites with p < 1×10^-5^. To determine whether mean autism effect sizes were significantly (p < 0.05) different between males and females, a Student’s two-tailed T-test was used to compare *Group* estimates from the within-cell-type, within-sex autism EWAS.

To identify cell-type-specific differences, the full list of cell type DMPs identified in any cell type from the within-cell-type EWAS analyses were evaluated in a model that utilises the entire dataset, with an additional covariate for cell type and an interaction term between cell type (*CellType*) and the main effect (i.e. *Group*CellType* or *Sex*Group*CellType*) (“across-cell-type EWAS”). The estimate for the main effect is with respect to the reference cell type of the *CellType* variable, which in this dataset was NeuN+ nuclei. The cell-type-by-main-effect interaction term splits into three interaction terms for each of the remaining non-NeuN+ cell types (SOX10+, IRF8+ and TN). For each DMP, the significance of each cell type was determined by both the significance of the main effect (p < or > 1×10^-5^) – which represents the significance of NeuN+ nuclei – and the interaction term (p > or < 0.05) – representing whether the estimate for the non-NeuN+ cell type is significantly different from that of the NeuN+ estimate. Incorporating data for all cell types increases the sensitivity to detect small effects consistent across cell types. Accordingly, DMPs originally detected in only one cell type from the within-cell-type EWAS may now have evidence for a significant effect in other cell types. Therefore, cell-type-specific DMPs were defined as sites that were significant only in one within-cell-type EWAS and had evidence for significance only in one cell type from the across-cell-type EWAS.

Across-cell-type EWAS model to test interaction of *Group* and *CellType*:

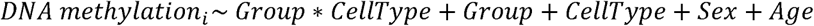

Across-cell-type EWAS model to test interaction of *Sex*Group* and *CellType*:

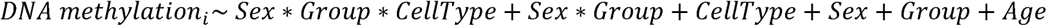

### Characterising sites with X chromosome inactivation status

X chromosome inactivation (XCI) status for 965 X-linked genes was obtained from San Roman *et al.* [81] (their Table S4A) and filtered to genes that are consensus “escape” from the cross-study meta-analysis. 93 of the 107 escape genes have at least one annotated site present in our dataset. X-linked DNA methylation sites were subsequently classified as “escape” if annotated to a consensus escape gene (2,062 sites annotated to 93 genes), and “non-escape” if annotated to any other X-linked gene (14,508 sites annotated to 955 genes). For each cell type, the distributions of the absolute within-female autism effect sizes between the two groups were compared statistically with a two-tailed T-test, with significant differences considered where p < 0.05.

### Comparison of DMP-associated genes with single-cell gene expression differences in autism

To determine whether genes associated with autism-related DNA methylation differences also show altered gene expression in autism, genes annotated to the 19 X-linked sex-by-autism DMPs identified in IRF8+ nuclei (**Table S4**) were interrogated using the ASD Single-Cell Gene Expression Portal [32] (available at http://solo.bmap.ucla.edu/asdscgene/). Differential expression results were examined for the MG1 and MG2 microglial cell populations with significant gene expression differences considered as those with FDR < 0.05.

### Co-methylation network analysis

We applied Weighted Gene Correlation Network Analysis (WGCNA) [85] to identify modules of co-methylated DNA methylation sites within each cell type. Prior to network construction, non-variable sites (sites whose range of the middle 80% of values is < 5% DNA methylation) and outlier values (where DNA methylation level > 3 standard deviations from the mean for that site) were excluded. WGCNA was performed separately for each cell type using a signed network with soft threshold power of 12 and minimum module size of 1,000 sites. Sites not assigned to a specific co-methylation module (the WGCNA “grey” module) were excluded from downstream analyses.

For each co-methylation module, the module eigengene (ME), defined as the first principal component of the DNA methylation level across sites assigned to the module, was regressed separately against i) autism diagnosis and ii) the interaction of diagnosis and sex, as described in the EWAS section earlier. Module membership (MM) for each site was defined as the correlation between its DNA methylation level and corresponding ME, with higher MM values indicating sites that are more representative of the overall module. Hub sites were defined as sites with an absolute MM > 0.7.

### Pathway analysis

Gene Ontology (GO) pathway analysis was performed separately for each autism-associated co-methylation module. GO terms were first limited to those containing between 10 and 2,000 genes (number of remaining terms = 7,542). For each GO term, logistic regression was used to test the association between gene set membership (binary variable of whether the gene is present in the GO term pathway) and module site count (number of module sites annotated to the gene), whilst controlling for gene size (total number of DNA methylation sites annotated to the gene). Significant GO terms were considered those with a z-distribution p-value associated with module status of less than 0.05 divided by the total number of terms tested (0.05 / 7,542 = 6.63×10^-6^). Significant GO terms were then collapsed to remove redundancy; the logistic regression model was repeated with each iteration controlling for genes present in the most significant term. To create the treemap plots for terms associated with the pink-IRF8+ module, the *calculateSimMatrix* and *reduceSimMatrix* functions from the *rrvgo* R package [127] were used with default parameters to calculate semantic similarity scores between GO terms and the *treemapPlot* function was used to plot the results.

## AVAILABILITY OF DATA

The dataset supporting the conclusions of this article is available in the Gene Expression Omnibus (GEO) repository, accession number GSE346075. Code relating to the analyses reported here can be found on GitHub (https://github.com/alicemfr/CellTypeAutismDNAm).

## AUTHOR CONTRIBUTIONS

Conceptualization: J.M., E.H., E.D. and A.F.; formal analysis: A.F., E.M.W., A.C.B; data generation: J.P.D., G.E.T.B., B.C., J.B.; supervision: J.M., E.H., R.B.; funding acquisition: J.M. and S.B-C; writing – original draft: A.F. and J.M.; writing – review & editing: all authors.

## SUPPLEMENTARY TABLES

**Table S1 - Distribution of samples within each nuclei fraction stratified by autism diagnosis and sex.** NeuN+ = neuron-enriched nuclei; SOX10+ = oligodendrocyte-enriched nuclei; IRF8+ = microglia-enriched nuclei; TN = triple negative, astrocyte-enriched nuclei (NeuN-/SOX10-/IRF8-). F = female (XX); M = male (XY).

**Table S2 – CETYGO-estimates of cell type composition within each FANS-isolated nuclei fraction.** Estimated proportions are shown for NeuN+, SOX10+, IRF8+ and triple-negative (NeuN−/SOX10−/IRF8−) nuclei. Two-tailed T-tests were used to compare mean estimated proportions between males and females and between autism and control groups.

**Table S3 - EWAS summary statistics for the 35 differentially methylated positions (DMPs) associated with autism.** For each cell type the estimated difference in mean DNA methylation level associated with autism (Estimate), its standard error (SE) and p-value (P) are reported. Columns starting “Sig” identify the cell type in which each DMP was detected. The chromosome (CHR) and genomic position (MAPINFO) are provided, alongside the gene annotation for sites within 1500bp upstream of a gene. Estimates and SEs are expressed as proportions of DNA methylation.

**Table S4 - EWAS summary statistics for the 59 differentially methylated positions (DMPs) associated with the interaction between autism and sex.** For each cell type the linear regression coefficient (Estimate), standard error (SE) and p-value (P) are reported for the sex-by-autism interaction (SexGroup) and the main effects of autism diagnostic group (Group) and sex (Sex). Columns starting “Sig” identify the cell type in which each DMP was detected. The chromosome (CHR) and genomic position (MAPINFO) are provided, alongside the gene annotation for sites within 1500bp upstream of a gene. Estimates and SEs are expressed as proportions of DNA methylation.

**Table S5 – Comparison of autism-associated effect sizes at autosomal sites between males and females.** Mean autism-associated effect sizes across 836,537 autosomal sites were calculated in males and females separately within each cell type. A T-test p-value of 0 indicates p < 1e-320. Effect sizes are expressed as proportions of DNA methylation.

**Table S6 – Comparison of autism-associated effect sizes at X-linked sites between males and females.** Mean autism-associated effect sizes across 20,609 X chromosome sites were calculated separately in males and females within each cell type. A T-test p-value of 0 indicates p < 1e-320. Effect sizes are expressed as proportions of DNA methylation.

**Table S7 – Comparison of autism-associated effect sizes in females according to XCI status.** Mean autism-associated effect sizes were compared between sites annotated to escape genes (2,062 sites annotated to 93 genes) or non-escape genes (genes normally subject to XCI) (14,508 sites annotated to 955 genes). SE = standard error of two-tailed T-test. Effect sizes are in units of percentage DNA methylation.

**Table S8 – Autism-associated gene expression differences in microglia for genes annotated to IRF8+ sex-by-autism DMPs.** For each of the 19 genes annotated to an X-linked IRF8+ sex-by-autism DMP, the autism-associated log fold change in gene expression (logFC) and FDR-corrected p-value (FDR) in MG1 and MG2 microglia are reported from the ASD Single-Cell Gene Expression Portal (Wamsley et al. 2024). The accompanying EWAS summary statistics in IRF8+ nuclei from the current study are provided for each DMP, including the p-value of the sex-by-autism interaction (IRF8.P.SexGroup), the difference in mean DNA methylation level associated with autism across both sexes (IRF8.Estimate.Group), in males only (IRF8.Estimate.Group.M) and in females only (IRF8.Estimate.Group.F) and the uncorrected p-values for these differences. The chromosome (CHR) and genomic position (MAPINFO) are provided, alongside the gene annotation for sites within 1500bp upstream of a gene. Alternative gene symbols (altGene) were used when the gene symbol in ‘Gene’ was absent from the Portal. NA indicates that expression data were unavailable from the Portal. Linear regression estimates are in units of percentage DNA methylation.

**Table S9 - Summary of WGCNA modules identified within each cell type.** WGCNA was performed on DNA methylation data subset to variable sites in each cell type. The total number of non-“grey” modules is shown. The first principal component of each module (module eigengene) was tested for its overall association with autism diagnosis and a sex-by-autism interaction, separately. Modules showing nominal significance (p < 0.05) for either association are indicated.

**Table S10 – Sites annotated to WGCNA modules in each cell type.** Non-variable sites that were excluded prior to WGCNA analysis are indicated by NA.

**Table S11 – Gene Ontology (GO) pathway-enrichment results for autism-associated WGCNA modules.** Presented are GO terms with a significant association between gene set membership and module site count (P.GenesinTestList < 6.63e-06, *logistic regression*) after collapsing to reduce redundant GO terms (indicated by MergeID and MergeName).

