## Supplementary Figures for "Cell-type-resolved DNA methylation profiling in the cortex reveals female-specific X chromosome differences in autism"

**Figure S1 – Generation of cell-type-resolved DNA methylation profiles using fluorescence-activated nuclei sorting (FANS). A)** Experimental workflow for generating genome-wide DNA methylation profiles from purified nuclei populations isolated from the human post-mortem prefrontal cortex (PFC). Bulk PFC tissue was homogenised to release nuclei, followed by fluorescence-based immunolabelling, FANS purification of discrete nuclei populations and DNA methylation profiling using the Illumina EPICv2 array. **B)** Representative FANS gating strategy used to isolate major cortical nuclei populations. Sequential gating was used to remove debris, doublets and non-nuclear events followed by separation of nuclei populations according to NeuN–Alexa Fluor 488, SOX10–NL577 and IRF8-APC fluorescence intensity. Representative plots show the identification of NeuN+ (neuronal) nuclei (purple), SOX10+ (oligodendrocyte) nuclei (dark pink) and IRF8+ (microglial) nuclei (blue) with NeuN-/SOX10-/IRF8- nuclei representing the remaining glial-enriched population (orange). Full experimental details are provided in Chioza *et al.* [1] and the accompanying protocol is available at https://dx.doi.org/10.17504/protocols.io.36wgq4965vk5/v5.

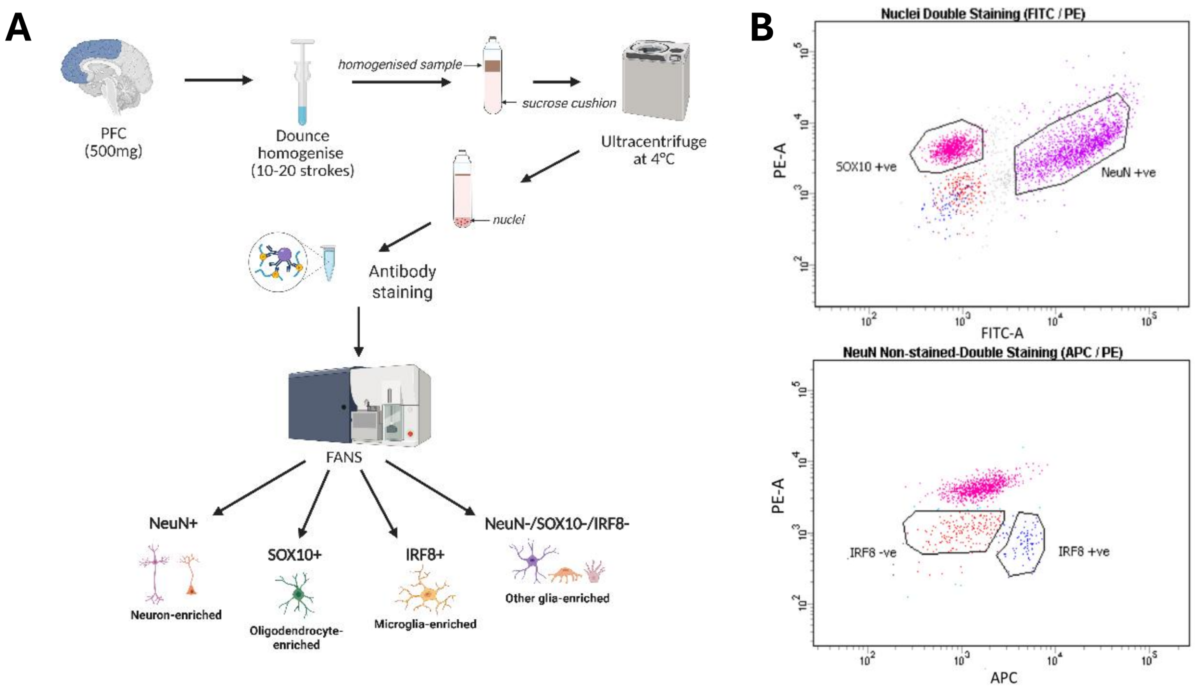

**Figure S2 – Demographic and sample characteristics of the study cohort. A-B)** Distribution of the 47 individuals included in this study stratified by sex and autism diagnosis. **C)** Distribution of samples from autistic and non-autistic donors across each purified nuclei population. **D-E)** Distribution of donor age stratified by sex and autism diagnosis. Dotted line indicates mean age (years).

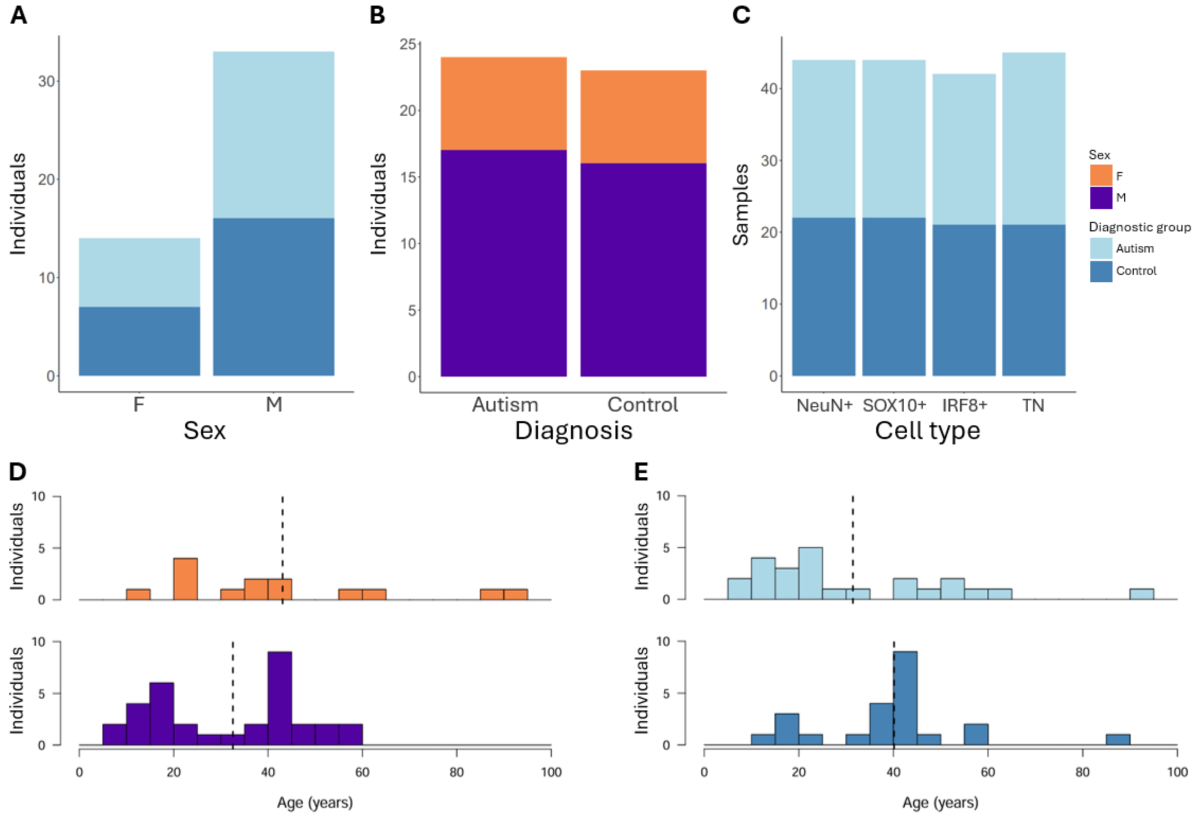

**Figure S3 – DNA methylation profiles cluster according to purified nuclei population.** Hierarchical clustering of the 10,000 most variable DNA methylation sites across all samples from neuron-enriched (NeuN+), oligodendrocyte-enriched (SOX10+), microglia-enriched (IRF8+) and astrocyte-enriched (TN; NeuN-/SOX10-/IRF8-) nuclei fractions isolated from prefrontal cortex. Samples cluster according to purified nuclei population, highlighting distinct DNA methylation profiles. DNA methylation proportions are represented from low (blue) to high (red).

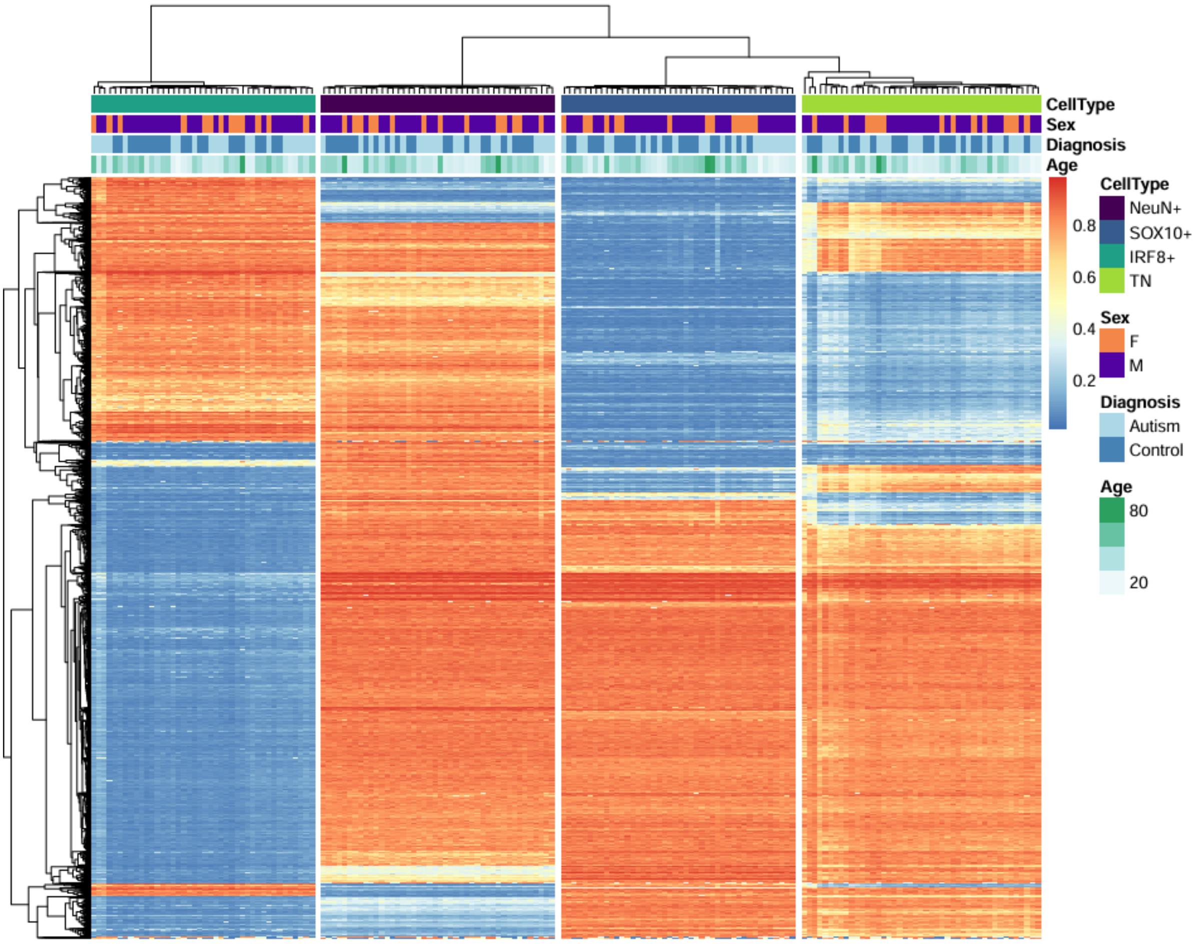

**Figure S4 – Single-nucleus RNA-sequencing confirms cellular identity of FANS-isolated nuclei populations.** Four FANS-isolated nuclei fractions from four post-mortem PFC samples were profiled using the Parse Biosciences Evercode low-input fixation workflow followed by single-nucleus RNA sequencing (see **Methods**). **A)** UMAP representation of nuclei coloured according to their FANS-isolated population (NeuN+, SOX10+, IRF8+ and TN [NeuN-/SOX10-/IRF8-]). **B)** UMAP representation coloured by predicted cell type identity following unsupervised clustering and annotation using the Seurat analysis workflow based on established cell type marker genes. **C)** Mean gene expression ($\log_e$(counts per 10,000 transcripts + 1)) of canonical marker genes for neurons (RBFOX3), oligodendrocytes (SOX10), microglia (P2RY12) and astrocytes (GFAP) across the four FANS-isolated nuclei populations.

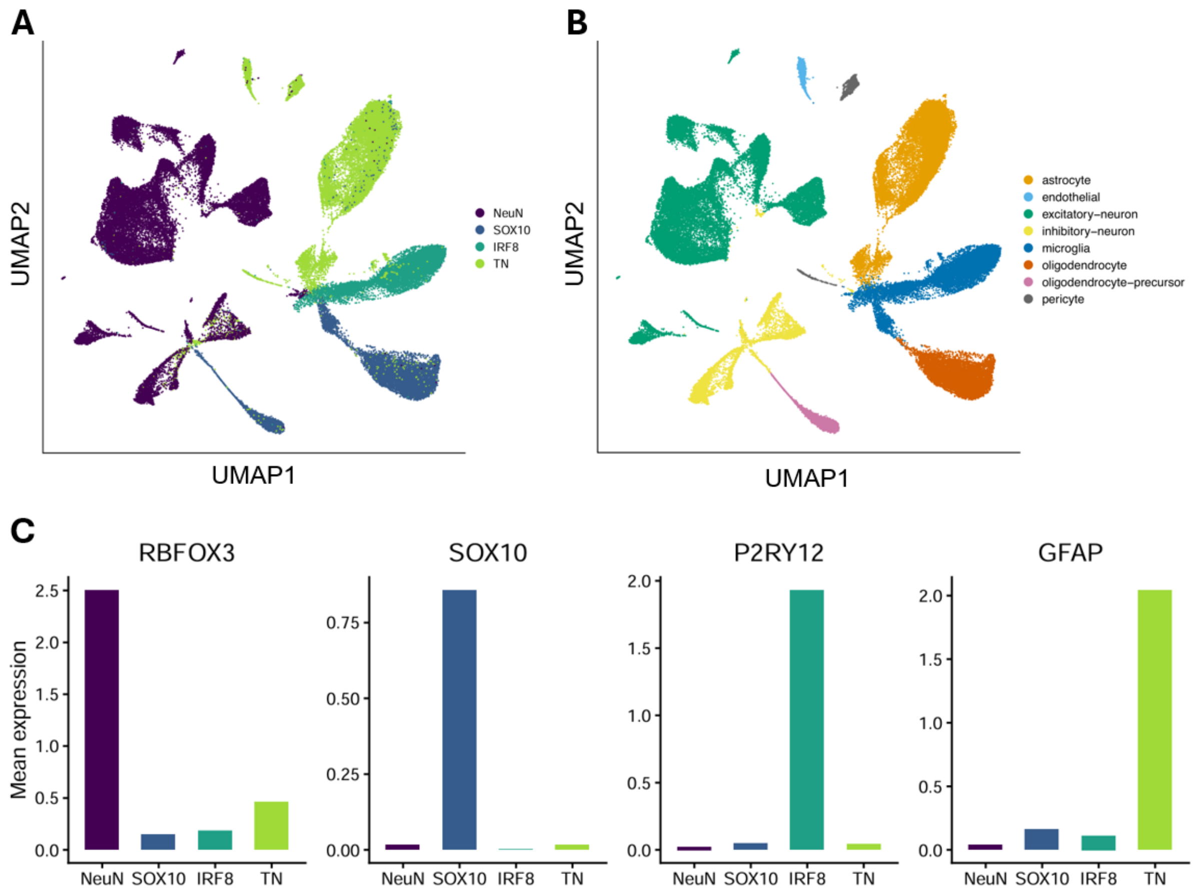

**Figure S5 – Confirmation of cell type enrichment of FANS-isolated nuclei populations.** **A)** CETYGO-estimated proportions of NeuN+, SOX10+, IRF8+ and NeuN-/SOX10-/IRF8- nuclei within each FANS-isolated nuclei population [2]. Plot titles indicate the FANS-isolated nuclei population for which cellular composition was estimated. **B)** CETYGO error metric for each cellular composition estimate. The red dashed line indicates an error metric of 0.1, with lower values indicating greater confidence in the estimated cell type proportions.

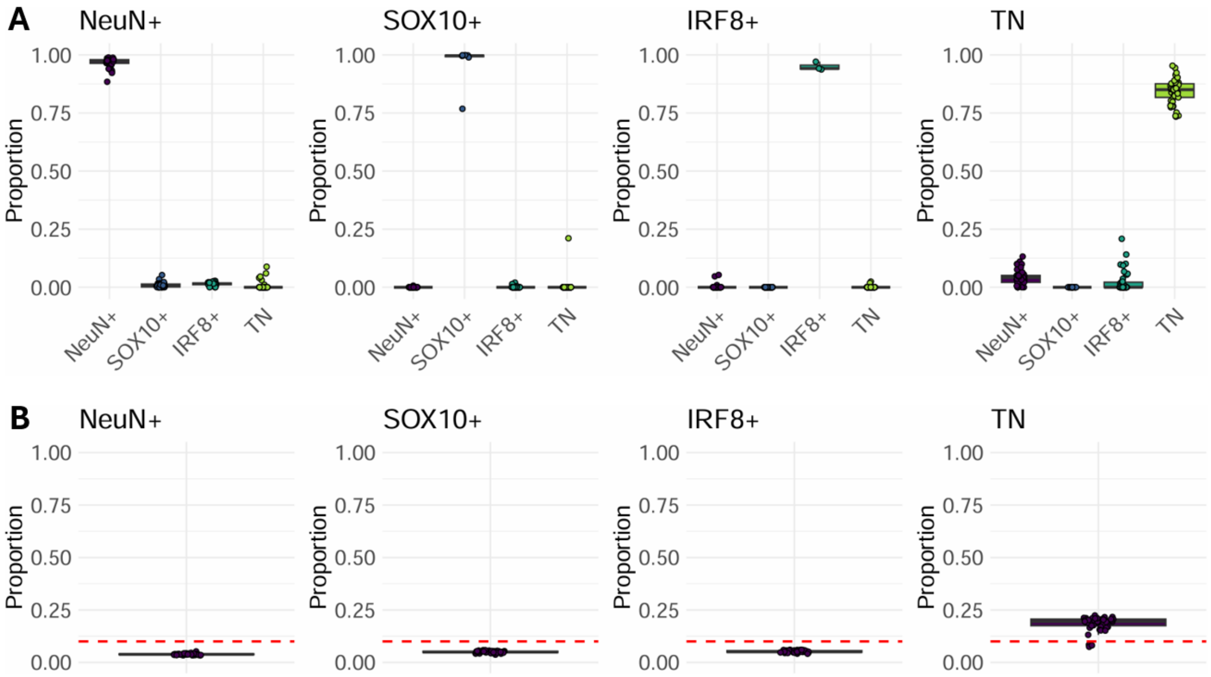

**Figure S6 – Genome-wide distribution of autism-associated DNA methylation differences across cortical cell types.** Manhattan plots showing the genomic distribution of -log10-transformed p-values from the autism EWAS performed separately in each purified nuclei population across the 857,146 DNA methylation sites tested. Sites reaching the discovery threshold ($p < 1x10^{-5}$) are labelled with their gene annotation, where available. The blue horizontal line indicates the discovery significance threshold of $p = 1x10^{-5}$.

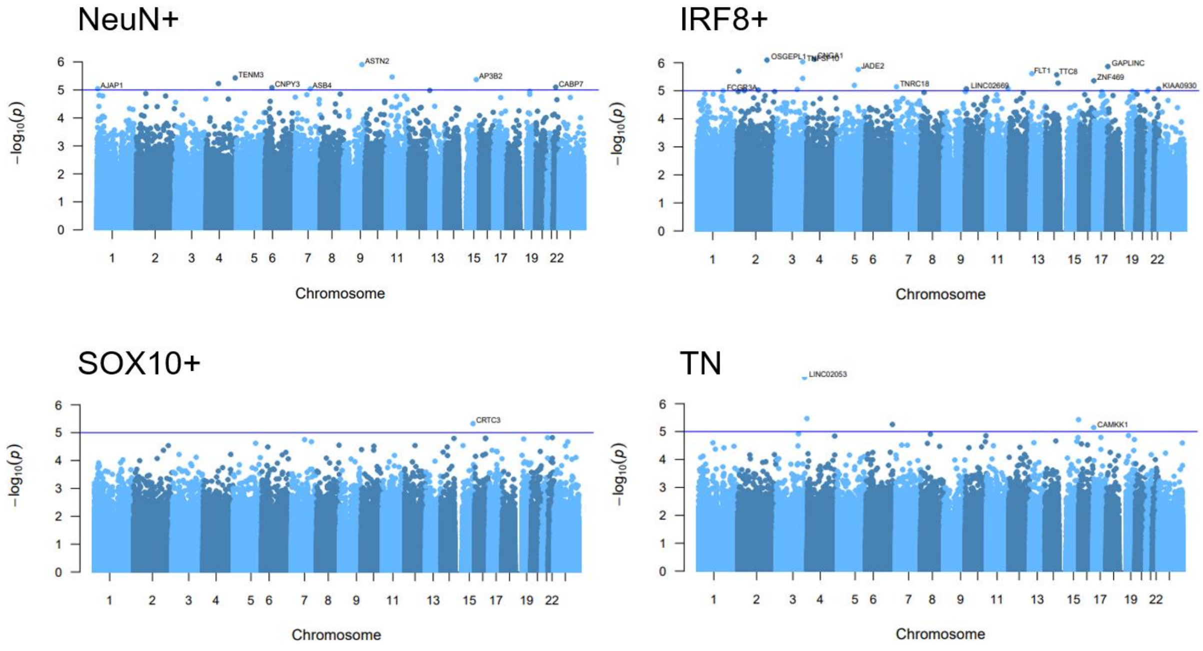

**Figure S7 – Representative autism-associated DNA methylation differences identified in each purified nuclei population.** Boxplots showing representative autism-associated DMPs identified in each purified nuclei population. Linear regression p-values are shown for cell types in which the autism association reached nominal significance ($p < 0.05$). **A)** cg08954634_TC21, annotated to *TENM3*, shows lower DNA methylation in NeuN+ neuronal nuclei from autistic individuals (mean autism-associated DNA methylation difference = 3.99%, p = $3.73 \times 10^{-6}$). *TENM3*, a member of the teneurin transmembrane protein family, has roles in neuronal migration [3] and synapse formation [4], two key features frequently dysregulated in autism. **B)** cg17830812_BC21, annotated to *CRTC3*, shows higher DNA methylation in SOX10+ nuclei from autistic individuals (mean autism-associated DNA methylation difference = -3.27%, p = $4.73 \times 10^{-6}$). *CRTC3* encodes a CREB-regulated transcriptional coactivator and is involved in cellular glucose metabolism, with disruption to energy metabolic processes being a notable feature in brains of autistic individuals [5]. **C)** cg03689904_TC21, annotated to *TMEM163*, shows lower DNA methylation in IRF8+ nuclei from autistic individuals (mean autism-associated DNA methylation difference = 4.02%, p = $9.39 \times 10^{-6}$). *TMEM163* has previously been implicated in immune and inflammatory pathways [6]. **D)** cg11979779_BC11, annotated to *CAMKK1*, shows higher DNA methylation in the astrocyte-enriched nuclei from autistic individuals (mean autism-associated DNA methylation difference = -1.68, p = $7.11 \times 10^{-6}$). *CAMKK1* encodes a component of the CaMK4 signalling pathway, which has been associated with autism [7].

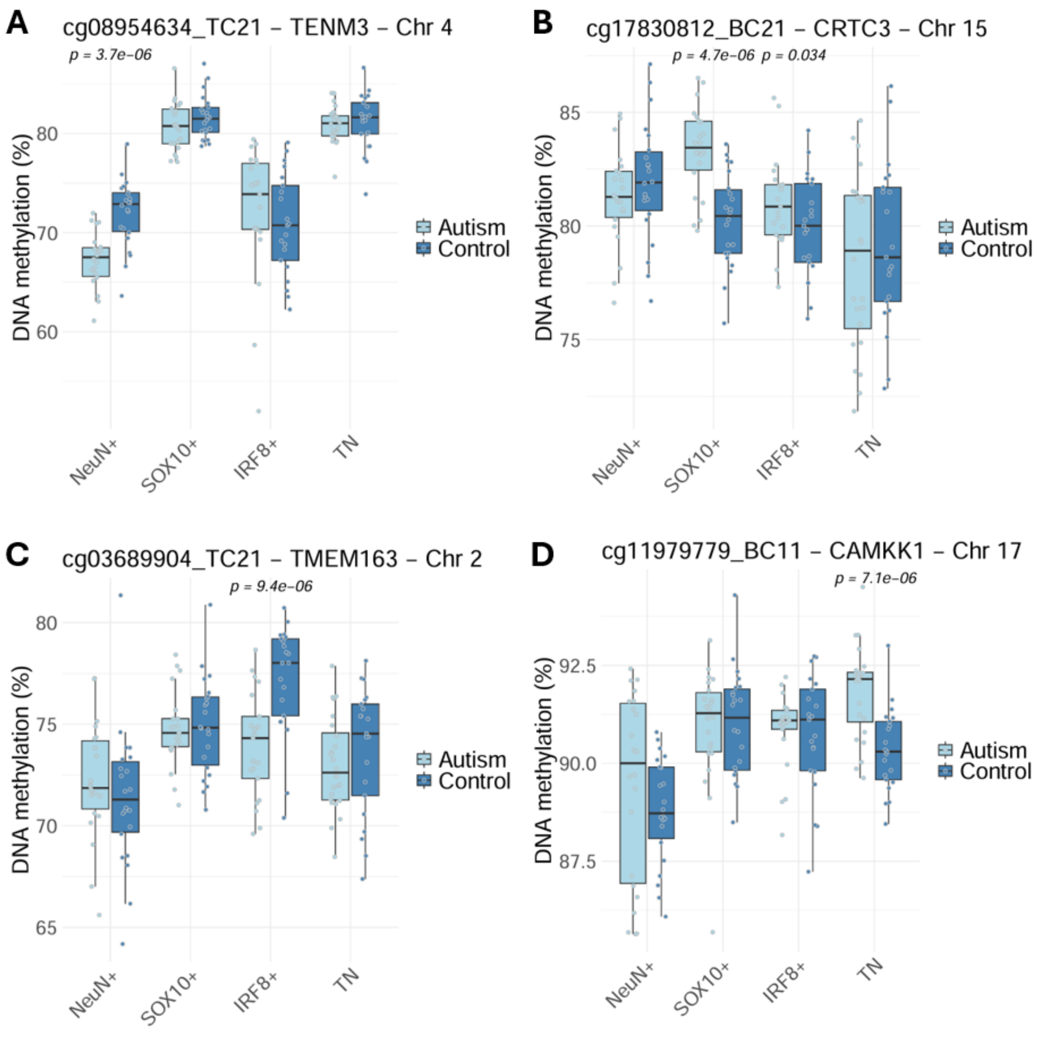

**Figure S8 – Correlations of autism-associated DNA methylation effect sizes between cell types are weakest for microglia.** Pairwise comparisons of autism-associated effect size estimates (mean autism-associated DNA methylation difference) across the 857,146 DNA methylation sites tested in each cell-type-specific EWAS. **A)** NeuN+ vs SOX10+ (r = 0.220), **B)** NeuN+ vs IRF8+ (r = 0.0655), **C)** NeuN+ vs TN (r = 0.203), **D)** SOX10+ vs IRF8+ (r = 0.0830), **E)** SOX10+ vs TN (r = 0.278), and **F)** IRF8+ vs TN (r = 0.0295). The solid line represents perfect correlation of effect sizes (y=x) and the dotted line represents the fitted linear regression line (y~x).

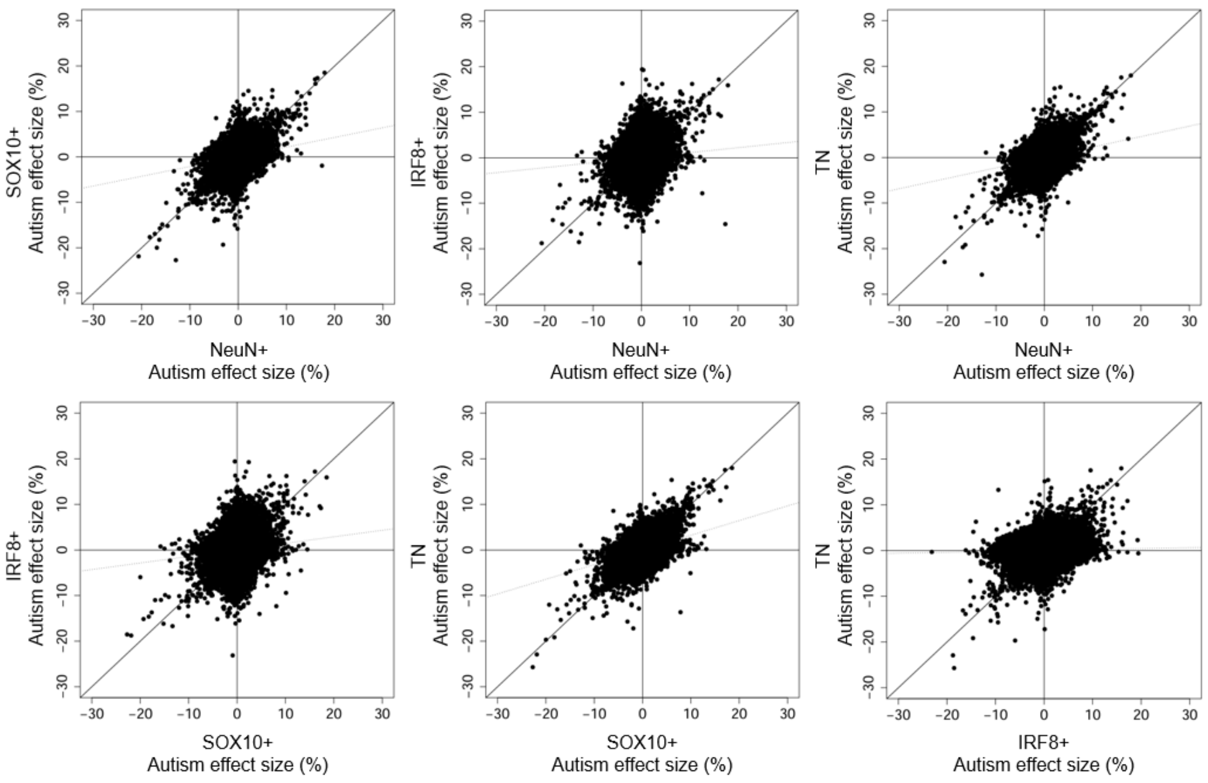

**Figure S9 – Autism-associated DNA methylation differences identified in bulk prefrontal cortex are most strongly correlated with microglial effects.** Comparison of autism-associated DNA methylation effect sizes identified in purified nuclei populations in the current study with those previously reported in bulk prefrontal cortex by [8]. Of the 31 autism-associated DMPs identified in the previous bulk cortex study, 28 were represented in the current dataset. Scatter plots show bulk cortex effect sizes plotted against the corresponding effect sizes in each purified nuclei population. Dotted lines represent fitted linear regression lines.

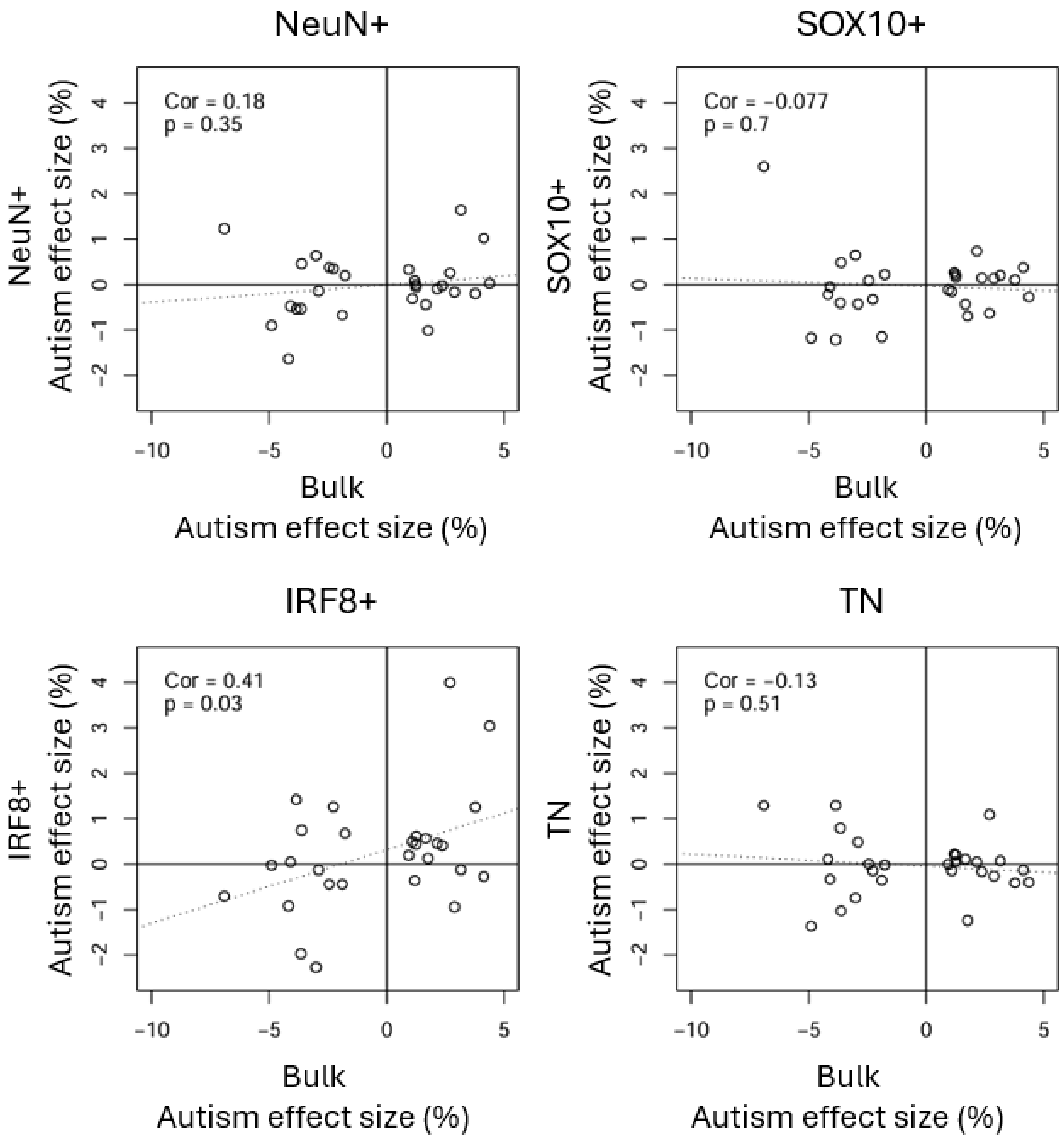

**Figure S10 – Sex-by-autism DMPs are annotated to genes with functional relevance to autism.** Boxplots showing representative sex-by-autism DMPs identified in each purified nuclei population. Sex-by-autism interaction p-values are shown for cell types in which the interaction reached nominal significance ($p < 0.05$). **A)** cg03752015_TC11 (sex-by-autism p = $5.12 \times 10^{-6}$), annotated to *CITED1*, demonstrates hypomethylation in neurons of autistic females (mean autism-associated DNA methylation difference = 5.08%, p = $1.85 \times 10^{-3}$) and no autism-associated differences in males (mean DNA methylation difference = -0.0640%, p = 0.860). *CITED1* is a regulator of estrogen-dependent transcription [9] and is required for placental development [10]. **B)** cg15039826_TC11 (sex-by-autism p = $6.01 \times 10^{-6}$), annotated to *BCOR*, exhibits hypomethylation in oligodendrocytes of autistic females (mean DNA methylation difference = 11.0%, p = $3.79 \times 10^{-3}$) and no significant difference in males (mean DNA methylation difference = -0.136, p = 0.747). *BCOR* regulates gene expression during early embryonic development [11] and a paralog of *BCOR* (*BCORL1*) has previously been associated with syndromic forms of autism [12]. **C)** cg18530240_BC21 (sex-by-autism p = $1.05 \times 10^{-7}$), annotated to *OGT*, demonstrates significant hypomethylation in microglia of autistic females (mean DNA methylation difference = 15.8%, p = $6.37 \times 10^{-4}$) and no significant difference in males (mean DNA methylation difference = -0.481, p = 0.663). *OGT* encodes the sole enzyme responsible for protein glycosylation and is a critical regulator of gene expression [13]. **D)** cg04875162_TC21 (sex-by-autism p = $3.97 \times 10^{-6}$), annotated to *PABPC5* – a regulator of polyadenylation and mRNA stability [14], exhibits hypomethylation in astrocytes of autistic females (mean DNA methylation difference = 5.79%, p = $1.32 \times 10^{-3}$) and no significant difference in males (mean DNA methylation difference = -0.405%, p = 0.427).

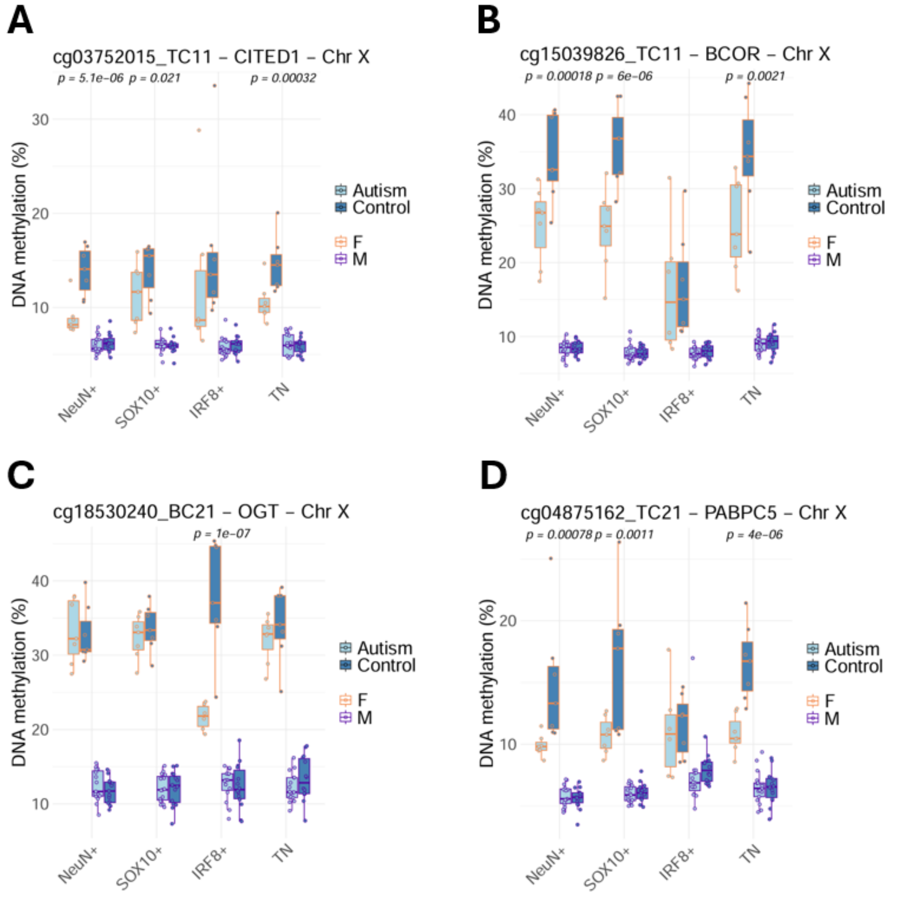

**Figure S11 – Genome-wide distribution of sex-by-autism interaction effects across cortical cell types.** Manhattan plots showing the genomic distribution of -log10-transformed p-values from the sex-by-autism EWAS performed separately in each purified nuclei population across the 857,146 DNA methylation sites tested. Sites reaching the discovery threshold ($p < 1x10^{-5}$) are labelled with their gene annotation, where available. Horizontal lines indicate significance thresholds ($p = 1x10^{-5}$ (blue), $p = 9x10^{-8}$ (red)).

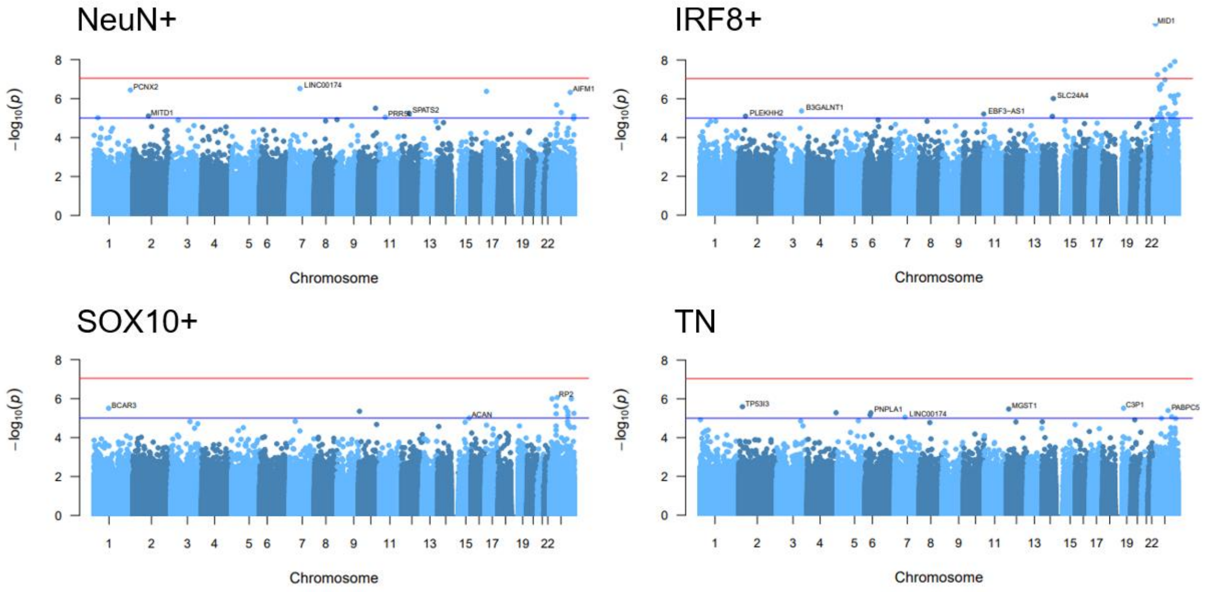

**Figure S12 – Autism-associated DNA methylation difference in the androgen receptor *(AR)* gene.** cg21966410 (sex-by-autism p = 1.66x10$^{-5}$), annotated to *AR*, shows hypomethylation in microglia of autistic females (mean autism-associated DNA methylation difference = 10.0%, p = 2.39x10$^{-3}$) and no significant difference in males (mean autism-associated DNA methylation difference = 0.113%, p = 0.817).

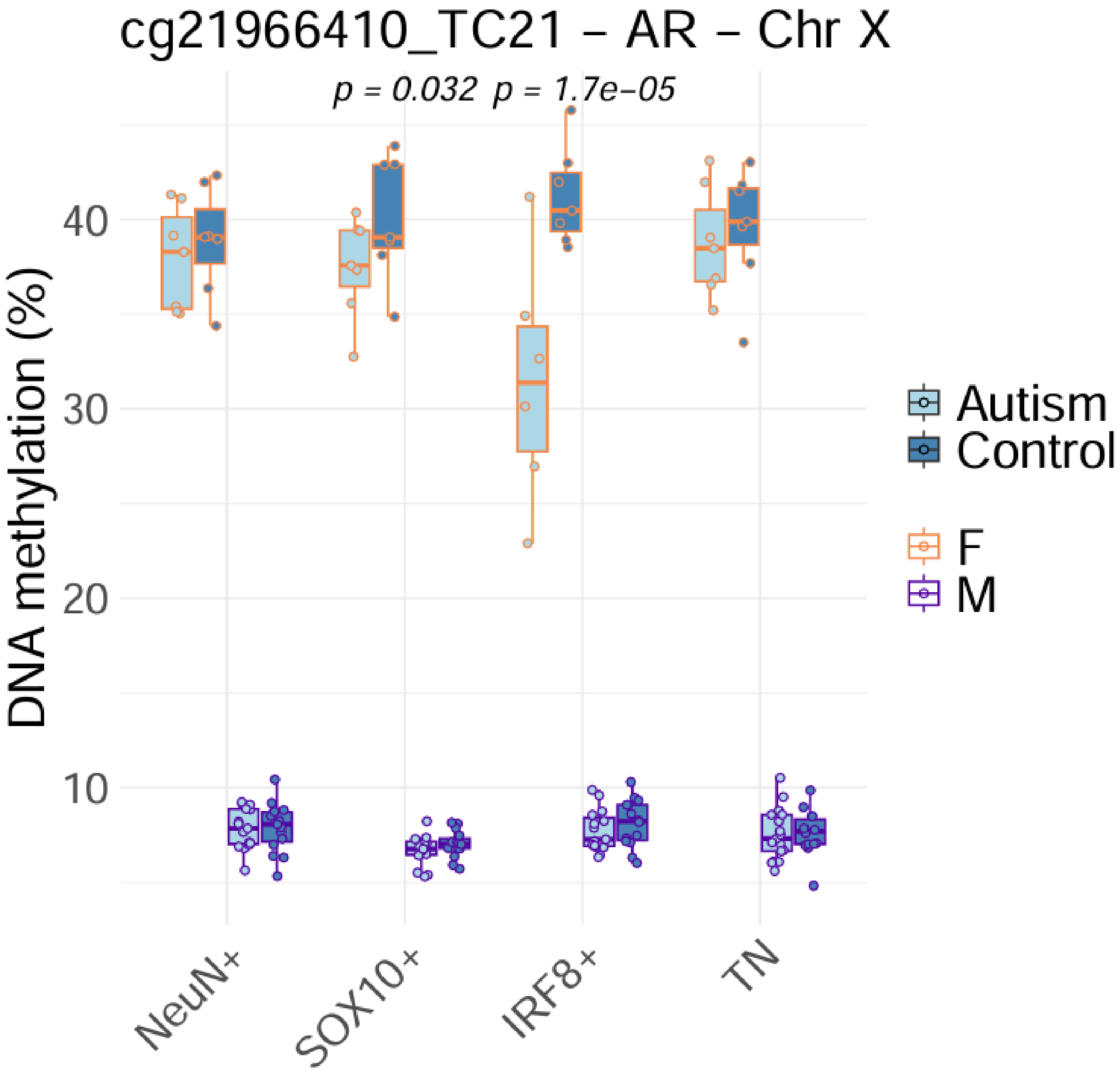

**Figure S13 – Autism-associated DNA methylation effect sizes are poorly correlated between males and females.** Comparison of autism-associated DNA methylation effect sizes estimated separately in males and females within each purified nuclei population. **A)** Autosomal sites (n = 836,537; correlation between male and female effect sizes: NeuN+ = 0.0499, SOX10+ = 0.0336, IRF8+ = 0.0537, TN = 0.0604). **B)** X-linked sites (n = 20,609; correlation between male and female effect sizes: NeuN+ = 0.0326, SOX10+ = 0.0271, IRF8+ = 0.0205, TN = 0.0213). Dotted line represents the fitted linear regression of female autism-associated effect size against the corresponding male effect sizes.

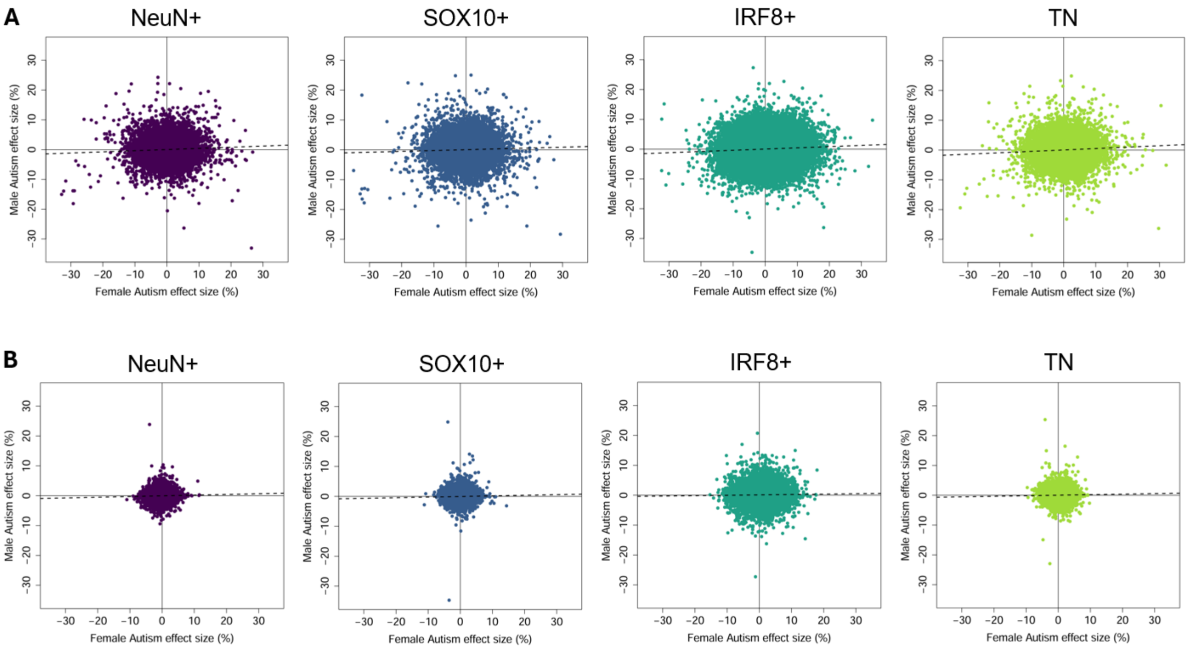

**Figure S14 – Module membership is associated with autism-associated DNA methylation effect size.** Scatter plots showing the relationship between module membership and effect size from the autism EWAS for the DNA methylation sites within each co-methylation module associated with autism. For sites in the magenta-IRF8+ module, module membership was regressed against the sex-by-autism EWAS effect size. Solid lines denote zero effect size (horizontal) and zero module membership (r = 0, vertical). Dashed line indicates regression of module membership against effect size.

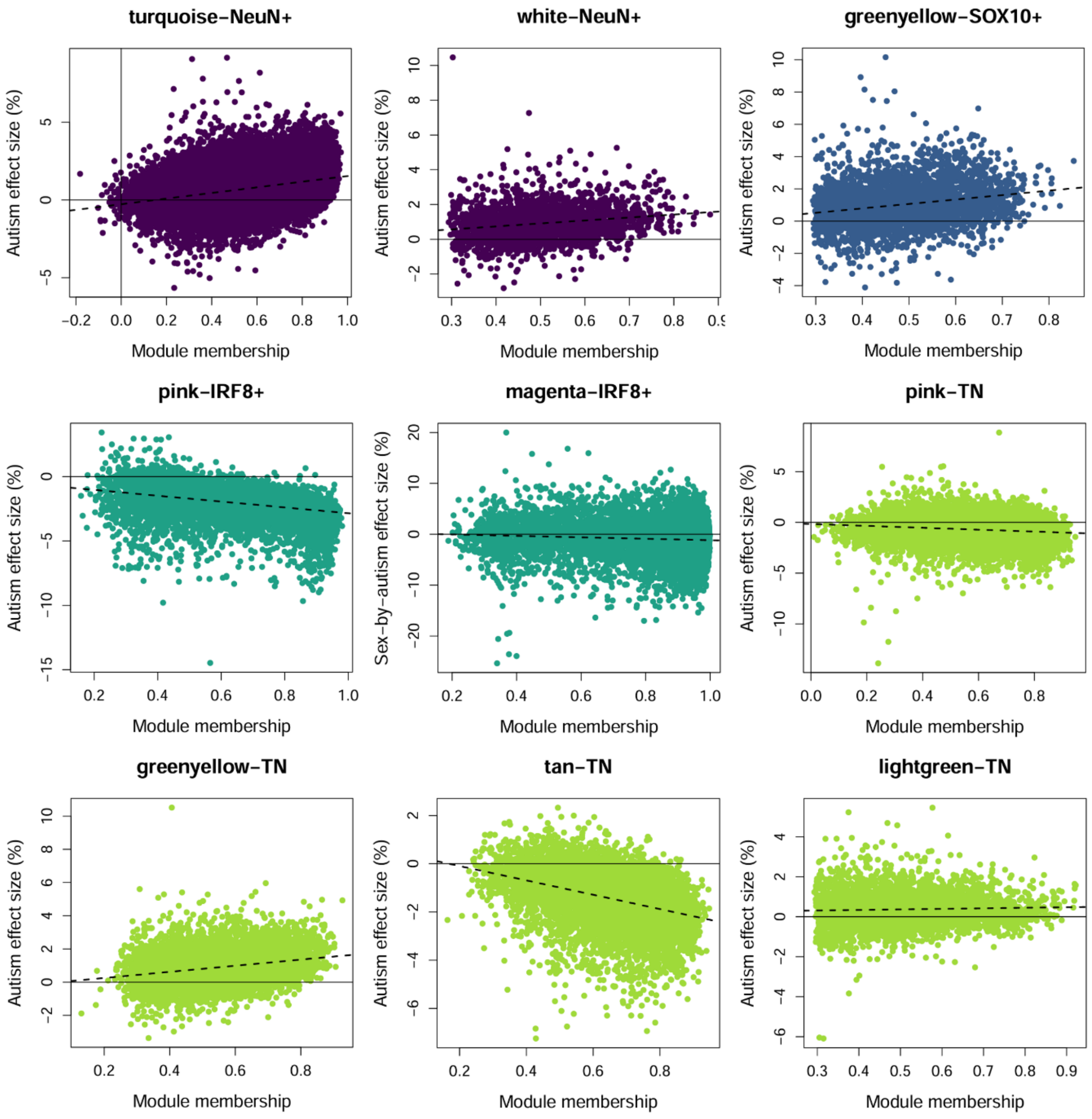

**Figure S15 – Hub sites within autism-associated co-methylation modules include genes implicated in autism and intellectual disability.** Boxplots showing representative hub sites from each autism-associated WGCNA co-methylation module. Many hub sites are annotated to genes previously implicated in autism and intellectual disability. Modules labelled **A-I**: turquoise-NeuN+, white-NeuN+, greenyellow-SOX10+, pink-IRF8+, magenta-IRF8+, pink-TN, greenyellow-TN, tan-TN, lightgreen-TN.

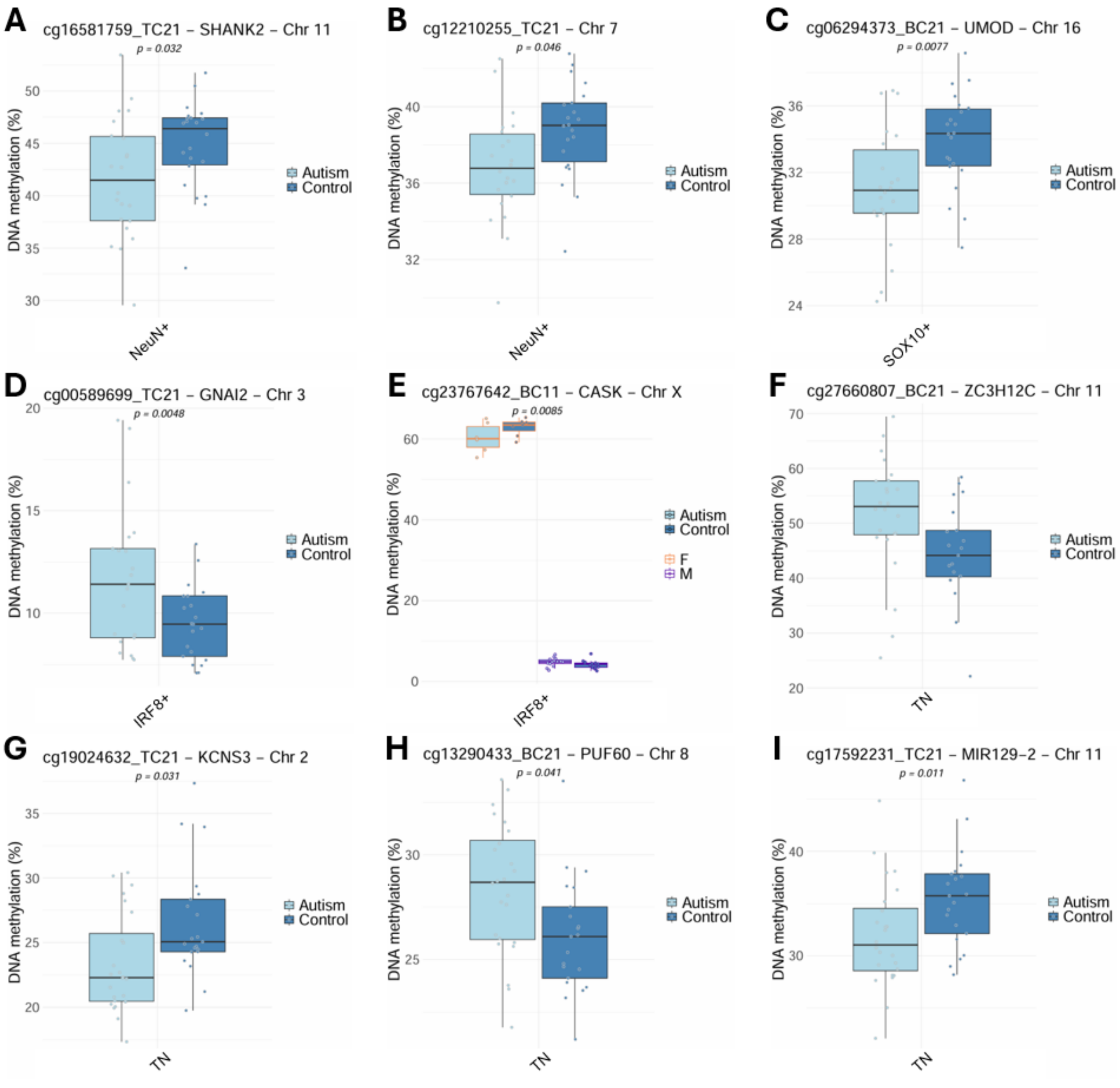

**Figure S16 – The sex-by-autism-associated microglial co-methylation module exhibits larger autism effect sizes in females.** Comparison of absolute autism effect sizes (mean autism-associated DNA methylation difference) between males and females at the 8,245 sites in the magenta-IRF8+ module. There is an overall significantly different mean absolute autism effect size between males (dotted line) and females (solid line) (difference in mean autism effect size = 1.56%, $p < 1x10^{-320}$, *two-tailed T-test*).

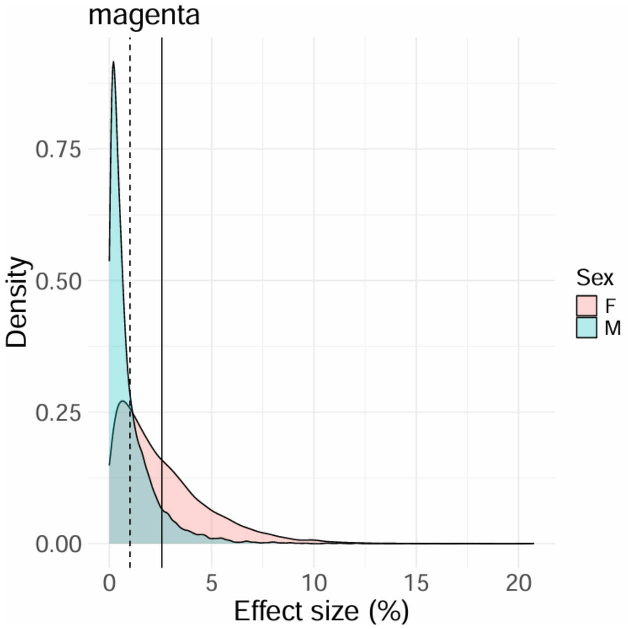

**Figure S17 – Autism-associated co-methylation modules are enriched for relevant biological pathways.** Presented are the top Gene Ontology (GO) terms enriched amongst genes annotated to sites in each module ($P < 6.63x10^{-6}$), including an enrichment for neurodevelopmental (turquoise-NeuN+), inflammatory (pink-IRF8+) and synaptic (magenta-IRF8+) pathways. Where more than 10 terms are significantly enriched, additional results can be found in **Table S11**.

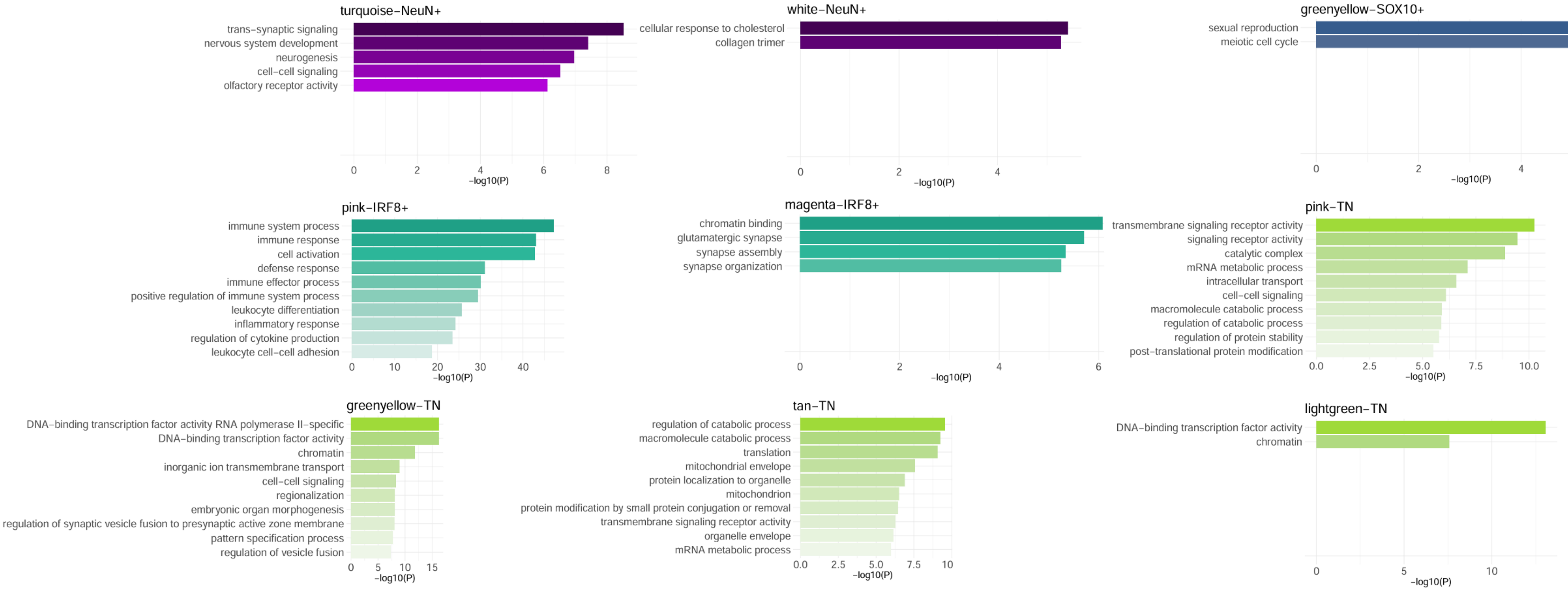

**Figure S18 – Enrichment of immune- and inflammatory-related pathways amongst genes annotated to sites in the pink-IRF8+ module.** Treemap plots showing Gene Ontology (GO) terms hierarchically clustered according to similarity scores (see **Methods**) for GO terms classified as **A)** biological process (BP) and **B)** molecular function (MF). Similar terms are grouped into blocks indicated by a thick outer border, with the highest scoring term (white text) as the representative term for the block.

**A**

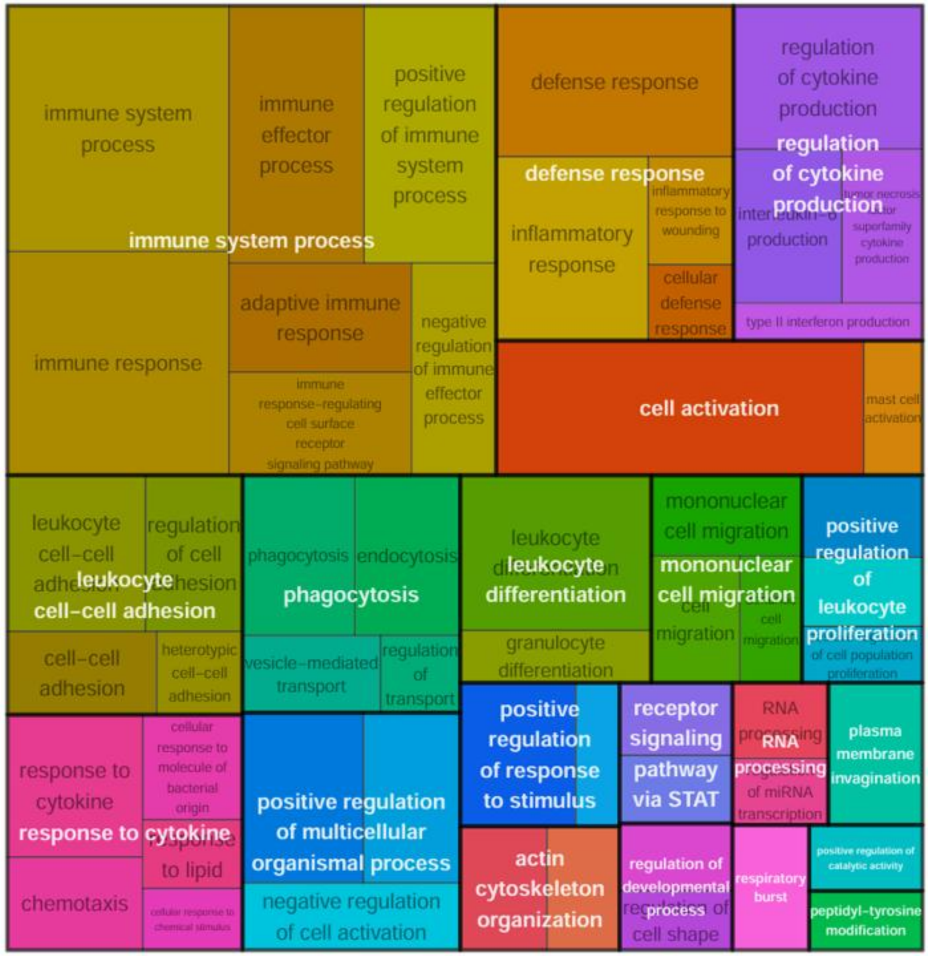


**B**

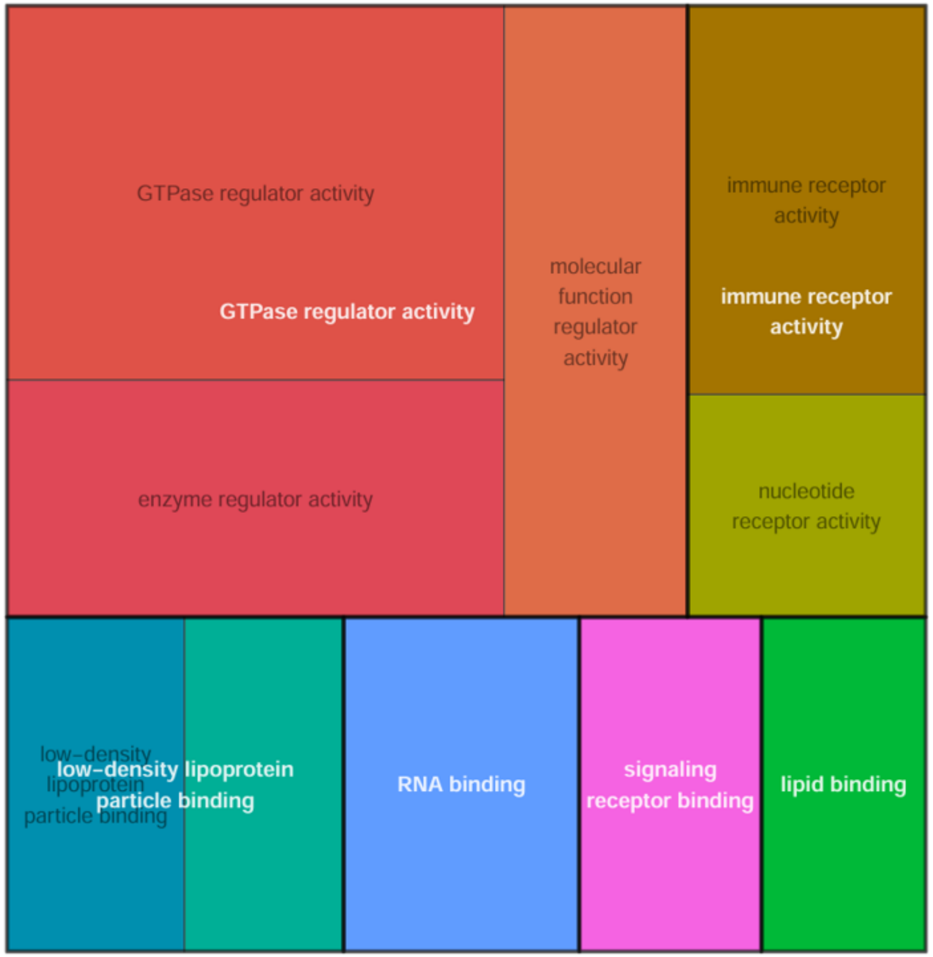

11. Wamstad JA, Corcoran CM, Keating AM, Bardwell VJ. Role of the Transcriptional Corepressor Bcor in Embryonic Stem Cell Differentiation and Early Embryonic Development.

PLoS One [Internet]. Public Library of Science; 2008 [cited 2026 Jul 17];3:e2814. https://doi.org/10.1371/JOURNAL.PONE.0002814

12. Shukla A, Girisha KM, Somashekar PH, Nampoothiri S, McClellan R, Vernon HJ. Variants in the transcriptional corepressor BCORL1 are associated with an X-linked disorder of intellectual disability, dysmorphic features, and behavioral abnormalities. Am J Med Genet A [Internet]. Wiley-Liss Inc.; 2019 [cited 2026 Jul 17];179:870–4. https://doi.org/10.1002/AJMG.A.61118

13. Lewis BA, Hanover JA. O-GlcNAc and the Epigenetic Regulation of Gene Expression. J Biol Chem [Internet]. American Society for Biochemistry and Molecular Biology Inc.; 2014 [cited 2026 Jun 9];289:34440. https://doi.org/10.1074/JBC.R114.595439

14. Blanco P, Sargent CA, Boucher CA, Howell G, Ross M, Affara NA. A novel poly(A)-binding protein gene (PABPC5) maps to an X-specific subinterval in the Xq21.3/Yp11.2 Homology block of the human sex chromosomes. Genomics [Internet]. Academic Press Inc.; 2001 [cited 2026 Aug 14];74:1–11. https://doi.org/10.1006/geno.2001.6530
